# A family of bacterial Josephin-like deubiquitinases with an unusual cleavage mode

**DOI:** 10.1101/2024.07.24.604956

**Authors:** Thomas Hermanns, Susanne Kolek, Matthias Uthoff, Richard A. de Heiden, Monique P.C. Mulder, Ulrich Baumann, Kay Hofmann

**Author notes:** Bayer AG, Research & Development, Pharmaceuticals, Biologics Research, Aprather Weg 18a, D-42113 Wuppertal, Germany. Correspondence should be addressed to Kay Hofmann.

## Abstract

Many intracellular bacteria secrete deubiquitinase (DUB) effectors into eukaryotic host cells to keep the bacterial surface or the enclosing vesicle membrane free of ubiquitin marks. Here, we describe a new family of bacterial DUBs that is structurally related to eukaryotic Josephins, but contains members that catalyze a unique destructive substrate deubiquitination. These ubiquitin C-terminal clippases (UCCs) cleave ubiquitin before the C-terminal diGly motif, thereby truncating the modifier and leaving a remnant on the substrate. By comparing the crystal structures of substrate-bound clippases and a closely related conventional DUB, we identified the factors causing the shift and found them conserved in other clippases, including one highly specific for M1-linked ubiquitin chains. This new enzyme class has great potential as tools to study the ubiquitin system, in particular aspects involving branched chains.

## Introduction

Protein ubiquitination is a posttranslational modification that regulates many aspects of eukaryotic cell biology, including the defense against intracellular pathogens. Through a cascade of ubiquitin activating, conjugating, and ligating enzymes, the C-terminus of ubiquitin is covalently attached to a substrate, usually via an isopeptide linkage to the ε-amino group of a substrate lysine, but occasionally to the N-terminus or to serine or threonine side chains^1^. Since substrate-attached ubiquitin can be ubiquitinated on several lysines, chains of different linkage types are generated, which confer different fates on the modified substrates. Ubiquitination can be reversed through the action of deubiquitinases (DUBs), which specifically cleave isopeptide or peptide bonds formed by the C-terminus of ubiquitin, thereby restoring both ubiquitin and the substrate to their original, unmodified state, with the possibility of later re-ubiquitination through the ubiquitin-conjugating cascade^2^.

Although bacteria lack a ubiquitin system on their own, many pathogenic or symbiotic bacteria have evolved ubiquitin-directed effectors that are secreted into the host cell and increase bacterial fitness by interfering with host defense pathways^3-5^. Bacterial ubiquitin ligases can modify host proteins with K48- or K11-linked ubiquitin chains, thereby targeting them for proteasomal degradation^6^. On the other hand, host cells use their own ligases to install ubiquitin on the surface of invading bacteria or on the surface of bacteria-containing vacuoles (BCVs), in which some intracellular bacteria are shielded from direct cytoplasmic access^7^. In the absence of bacterial countermeasures, this surface-bound ubiquitin targets bacterial particles for xenophagy or directs ubiquitinated BCVs toward lysosomal degradation^8^. Bacteria with intracellular lifestyles have evolved mechanism to evade this fate, either by preventing ligase access to the bacteria^9^, by interfering with ubiquitination^10^, or – most prominently – by using DUB effectors to remove previously attached ubiquitin ^3, 11, 12^. Most bacterial DUBs appear to be recent acquisitions from host genomes, since they show recognizable sequence and structural similarity to eukaryotic DUBs, and are often restricted to narrow bacterial taxa. More distantly related bacteria often encode DUBs that result from independent acquisition events. Apart from a small metalloprotease family, all eukaryotic DUBs are papain-fold cysteine proteases, which can be grouped into seven different classes (USP, UCH, OTU, Josephin, MINDY, ZUFSP, and VTD) ^2, 13^. Most bacterial DUBs are either related to the OTU (ovarian tumor) family ^14^ or belong to the so-called CE-clan, an enzyme family comprising eukaryotic proteases for the ubiquitin-like modifiers SUMO and NEDD8 and bacterial deubiquitinating enzymes ^15^. Typical intracellular bacteria such as *Salmonella typhimurium*, *Shigella flexneri*, *Burkholderia pseudomallei*, *Chlamydia pneumoniae*, or *Chlamydia trachomatis* encode one or two deubiquitinases, usually members of the OTU or CE family with little linkage specificity ^11^. Two exceptions are *Legionella pneumophila* and *Simkania negevensis*, two unrelated bacteria with a wide host range that encode a large and diverse set of deubiquitinases, some of which are highly specific for K6-linked or linear chains ^16-18^.

Recently, Simkania was found to encode two members of the Josephin family, a somewhat enigmatic DUB class previously thought to be eukaryote-specific ^18^. During the characterization of SnJOS1 and SnJOS2, we made the surprising observation that these two enzymes show an unusual cleavage mode: When incubated with ubiquitin chains, both SnJOS1 and SnJOS2 did not cleave the isopeptide bond behind the ubiquitin C-terminus, but rather the peptide bond between Arg-74 and Gly-75 of ubiquitin. The same bond is also cleaved in mono-ubiquitin, resulting in a non-functional ubiquitin C-terminally shortened by two residues. This activity, which we refer to as ubiquitin C-terminal clippase (UCC), was also observed in many, but not all, additional bacterial Josephin relatives from several bacterial phyla. Their reaction amounts to destructive deubiquitination, since the shortened ubiquitin cannot be re-conjugated, and the diGly remnant left on the substrate lysine precludes further modification of this residue. Eukaryotic Josephins, however, cleave ubiquitin at the canonical DUB position behind Gly-76. By solving the substrate-bound structures of a linkage-promiscuous UCC from *Burkholderia pyrrocinia* (BpJOS) and the closely related conventional DUB from *Pigmentiphaga aceris* (PaJOS), we determined the structural basis of the discordant cleavage positions. Some bacterial UCCs were found to be linkage-specific: the UCC from *Parachlamydia sp*. (PcJOS) cleaved only linear ubiquitin chains, a specificity that could be rationalized through analysis of the PcJOS structure in complex with linear diubiquitin. Taken together, bacterial clippases of the Josephin family are the first enzymes to show this activity, which allows the bacteria to make deubiquitination an irreversible process.

## Results

### Simkania Josephins are ubiquitin C-terminal clippases

While characterizing the first bacterial Josephin-type DUBs from *Simkania negevensis*, we found them to be inactive against the mono-ubiquitin-based model substrates Ub-AMC and Ub-PA, whereas both enzymes were similarly active against diubiquitin species of different linkage types ^18^. Since linkage-promiscuous DUBs usually do react with AMC- or PA-based model substrates, we further investigated the chain cleavage mode. Despite its reactivity against K63-liked diubiquitin, SnJOS1 failed to react with a K63-linked diUb-VME probe (Supplementary Fig 1a), suggesting that the lack of Ub-AMC and Ub-PA reactivity is not due to the absence of the proximal (S1’) ubiquitin moiety. While analyzing the products of K63 diubiquitin cleavage, we observed cleavage products with masses of ±114 Da, in addition to the expected mono-ubiquitin (Fig 1a). This mass difference corresponds to the diGly peptide found at the C-terminus of ubiquitin, suggesting a cleavage after Arg-74 of both ubiquitin units. A similar reaction was observed when incubating SnJOS1 or SnJOS2 with mono-ubiquitin, which resulted in the loss of the terminal diGly (Δ114 Da) from the substrate (Supplementary Fig 1b). Such an activity, which we call ‘ubiquitin C-terminal clippase’ (UCC), has never been observed in physiological enzymes but is reminiscent of the ISG15-shortening activity of the picornaviral leader peptidase Lb^pro 19^, which has been engineered to also act on ubiquitin ^20^. The diGly remnant left on the proximal cleavage product can be detected by a Lys-ε-Gly-Gly antibody as shown in Fig 1b, which monitors the degradation of K63-linked Ub_6+_ by SnJOS1/2. Both Simkania clippases show a time-dependent shortening of the input chains with concomitant accumulation of Lys-ε-Gly-Gly on the proximal ubiquitin. In the case of SnJOS1, only diGly modified mono-ubiquitin was left at 6h, while the reaction progress of the slower SnJOS2 at 6h resembles the 1h timepoint of SnJOS1. To further confirm the cleavage position after Arg-74 and address the question of whether the lack of reactivity against Ub-PA is due to the unconventional cleavage site, we generated a shortened activity-based probe (Ub^1-73^-PA) in which Arg-74 was replaced by the reactive propargylamine (PA) warhead ^21^. Indeed, both SnJOS1 and SnJOS2 reacted exclusively with Ub^1-73^-PA, but not with the conventional probe Ub^1-75^-PA (Fig 1c), demonstrating that these enzymes do react with activity-based probes but require a reactive group at the correct position.

**Figure 1:**
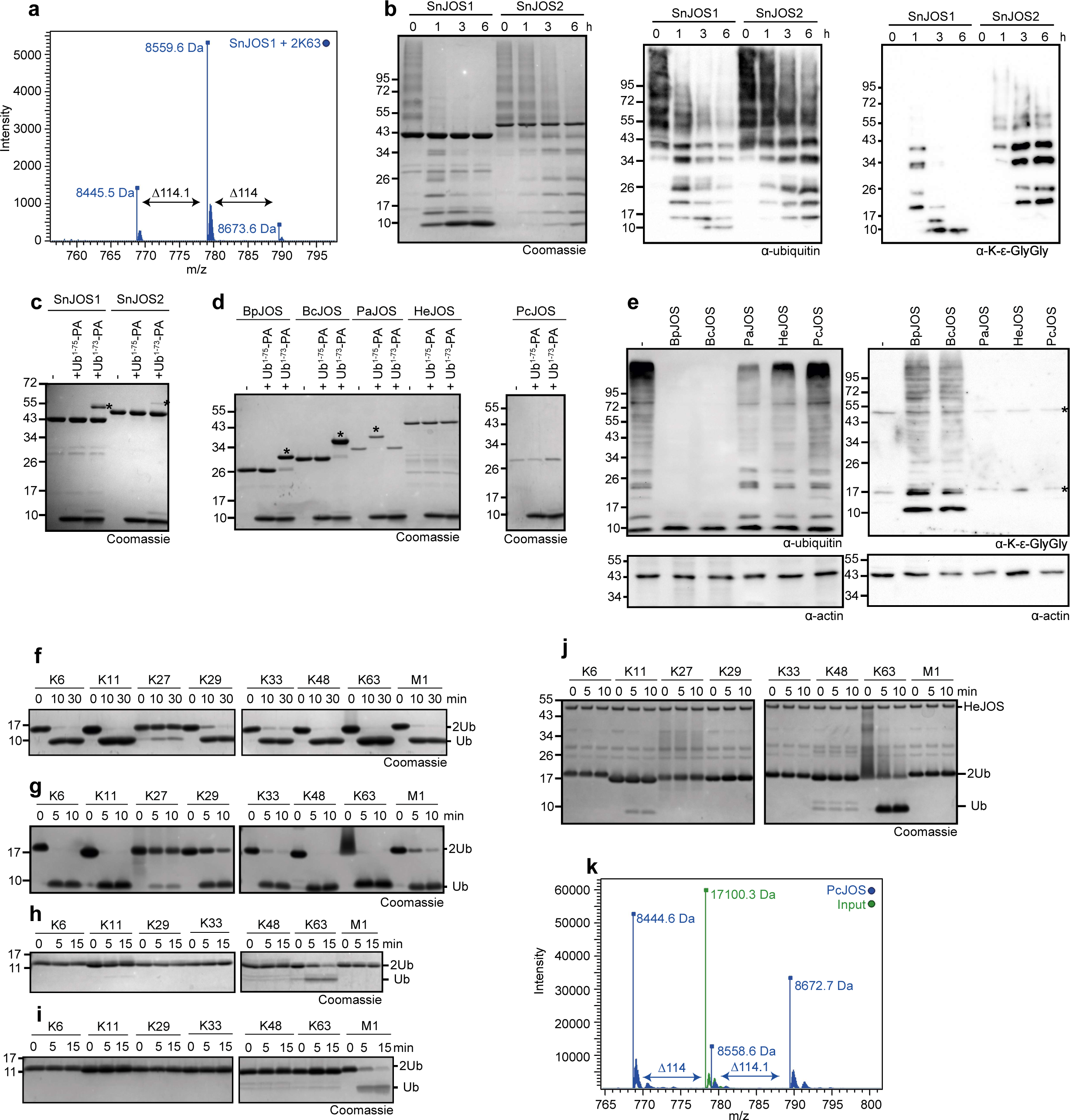
Bacterial Josephins cleave ubiquitin at different positions. **a)** Intact mass spectrometry of K63-linked diubiquitin cleaved by SnJOS1. The m/z ratio is depicted on the x-axis, while the deconvoluted masses are shown next to the respective peaks. The 8559.6 Da peak corresponds to the monoisotopic mass of mono-ubiquitin, whereas the remaining peaks differ by 114 Da indicating the removal/addition of a GlyGly-peptide. **b)** Analysis of SnJOS1/2 cleavage products by Western Blotting. K63-linked Ub_6+_ chains were incubated with 5 µM SnJOS1 or SnJOS2 for the indicated time points. Time-dependent degradation of the poly-ubiquitin chains is visualized by Coomassie-staining (left panel) and α-ubiquitin western blotting. Simultaneous accumulation of GlyGly-remnants is visualized by α-K-ε-GlyGly detection (right panel). **c)** Determination of the cleavage position by activity-based probes. SnJOS1 and SnJOS2 were incubated with Ub^1-75^ or Ub^1-73^-PA probes for 18h. Asterisks (*) mark the shifted bands after the reaction. **d)** Differential probe reactivity of additional bacterial Josephin homologues, performed according to c). **e)** Analysis of bacterial Josephins cleavage products by Western Blotting. HEK293T cell lysates were incubated with the BpJOS, BcJOS, PaJOS, HeJOS or PcJOS for 1h. Substrate deubiquitination and accumulation of mono-ubiquitin is visualized by α-ubiquitin detection (left panel) and accumulation of diGly-remnants is visualized by α-K-ε-GlyGly detection (right panel). α-actin staining serves as a loading control. Asterisks (*) mark unspecific bands. **f-j)** Linkage specificity analysis of bacterial Josephin-DUBs. A panel of diubiquitin chains was treated with 50 nM BpJOS (f), 50 nM BcJOS (g), 50 nM PaJOS (h), 0.25 µM PcJOS (i) or 2.5 µM HeJOS (j) for the indicated time points. **k)** Intact mass spectrometry of M1-linked diubiquitin cleaved by PcJOS. The m/z ratio is depicted on the x-axis, while the deconvoluted masses are shown next to the respective peaks. The input sample shows a single 17100 Da peak (green) corresponding to the monoisotopic mass of diubiquitin, which gets cleaved by PcJOS to three different products (blue) corresponding to the mono-isotopic mass of ubiquitin (8558.6) or ubiquitin ± diGly peptide (8672 / 8444 Da).

### A bacterial Josephin family with different cleavage activities

Eukaryotic Josephins have been consistently described as conventional deubiquitinases^22-24^. We verified the published results using our assays and found that human ATXN3L and JOSD2 lack clippase activity (Supplementary Fig 2). To investigate whether SnJOS1/2 are exceptional cases, or if the shifted cleavage position is a hallmark of a bacterial UCC family, we performed bioinformatical database searches for additional bacterial Josephin homologs ^25^. Using generalized profile searches ^26^, we identified related sequences in the genomes of several bacteria, including *Burkholderia pyrrocinia* (BpJOS), *Burkholderia catarinensis* (BcJOS), *Pigmentiphaga aceris* (PaJOS), *Herbaspirillum sp*. (HeJOS) and *Parachlamydia sp.* (PcJOS) (Supplementary Fig. 3). We expressed the predicted catalytic domains of these enzymes in *E. coli*, purified them, and characterized their catalytic properties. The *Burkholderia* Josephins BpJOS and BcJOS reacted only with the truncated Ub^1-73^-PA probe, suggesting a clippase activity. By contrast, *Pigmentiphaga* PaJOS reacted exclusively with the conventional Ub^1-75^-PA probe, indicating that UCC activity is not a general feature of the bacterial Josephin family (Fig 1d). The cleavage positions were confirmed by intact mass spectrometry (Supplementary Fig 1c-e). The two remaining candidates HeJOS and PcJOS reacted with neither probe, suggesting that they are inactive or require a particular linkage type for their activity (Fig 1d). When incubating the purified enzymes with HEK293 cell lysate and visualizing the products by ubiquitin- or Lys-ε-diGly-directed antibodies, the results supported the probe reactivity data (Fig 1e). BpJOS and BcJOS completely deconjugated all cellular ubiquitin and left diGly remnants on their substrates. In line with the probe results, PaJOS did not generate diGly remnants, although some ubiquitin was de-conjugated. HeJOS appeared similar to PaJOS, with no diGly production but a modest reduction in high-MW chains. PcJOS appeared inactive in this experiment, as neither chain reduction nor antibody-detectable diGly remnants were observed (Fig 1e).

Linkage specificity was analyzed using a panel of differently linked diubiquitins. The Burkholderia clippases BpJOS and BcJOS cleaved all chain types within minutes, with the exception of K27-linked diubiquitin, which was only poorly cleaved (Fig 1f,g). By contrast, the conventional DUB PaJOS was highly specific for K63-linked chains, highlighting the differences within the bacterial JOS family (Fig 1h). HeJOS and PcJOS, which were inactive against activity-based probes, specifically cleaved K63- or M1-linked chains, respectively (Fig 1i,j). To determine the cleavage position of the linkage-specific Josephins, the proteases were incubated with the respective diubiquitin species, and the generated mono-Ub was analyzed using intact MS. The HeJOS-generated ubiquitin had a mass of 8558.6 Da, indicating conventional DUB cleavage (Supplementary Fig 1f). In case of PcJOS, the cleaved linear diubiquitin showed masses of 8444.6 Da, 8558.6 Da and 8672.6 Da, indicating that the C-terminal diGly was cleaved off the distal ubiquitin and remained attached to the N-terminus of the proximal ubiquitin (Fig 1k). The free C-terminus of the proximal ubiquitin was apparently poorly clipped.

The activity of BpJOS as a linkage-promiscuous clippase is reminiscent of Lb^pro^*, a variant of the viral leader peptidase Lb^pro^ engineered to also work on ubiquitin chains, which are cleaved without linkage selectivity ^20^. We compared the activity of BpJOS and Lb^pro^ towards ubiquitin and ubiquitin-like modifiers. While Lb^pro^* reacted with clippase probes Ub^1-73^-PA, NEDD8^1-73^-PA, and ISG15^79-154^-PA, BpJOS only reacted with the ubiquitin and NEDD8 probes (Supplementary Fig 4a). The inactivity of BpJOS towards ISG15 was confirmed by intact mass spectrometry (Supplementary Fig 4b). By contrast, the activity of BpJOS towards K48-linked diubiquitin exceeded that of Lb^pro^* by several orders of magnitude (Supplementary Fig 4c).

### Structural basis of M1-specific ubiquitin clipping by PcJOS

To understand the structural basis of the shifted UCC cleavage site in a linkage-specific context, we determined the structure of PcJOS in complex with its substrate. The catalytic fragment PcJOS^70-324^, rendered inactive through mutation of the predicted active site residue Cys-162 to alanine (Supplementary Fig 3), was crystallized with linear diubiquitin and the crystal structure solved to a resolution of 2.18 Å. The asymmetric unit contained one diubiquitin molecule bound by two PcJOS molecules, the first bound between the ubiquitin units, and the second bound to the C-terminus of the diubiquitin (Fig S5a). In the first PcJOS^70-324^ molecule (chain A), region 97-324 was fully resolved; chain C showed a nearly identical conformation with an RMSD of 0.44 Å over 1317 atoms (Fig S5b). The PcJOS catalytic domain comprises a papain-fold core domain reminiscent of other Josephin-type DUBs (α1-β6) and an N-terminal extension formed by helices α1’-α3’ (Fig 2a). The active site consists of Cys-162 (Ala-162 in the structure), His-284, and Asp-300, in line with the alignment-based prediction (Supplementary Fig 3). In the first PcJOS molecule, the active site is placed next to the scissile peptide bond between Arg-74 and Gly-75 of the distal (S1) ubiquitin, compatible with the observed clippase activity (Fig 2b). Mutating any active site residue to alanine caused a complete loss of cleavage activity (Supplementary Fig 5c). Structural database searches using DALI^27^ yielded several Josephin DUBs as the best matches, including human JOSD2 (PDB:6PGV)^24^ and ATXN3L (PDB:3O65)^22^, thus supporting the assignment of PcJOS to the Josephin family. The structural similarity was most pronounced within the catalytic core region (Supplementary Fig 5d) including the active site residues (Supplementary Fig 5e); structural superposition of the core region resulted in RMSDs of 1.4 Å over 500 atoms for JOSD2 and 3.7 Å over 574 atoms for ATXN3L. Two striking differences between PcJOS and eukaryotic Josephins are the extended N-terminal helix bundle (α1’-α3’, residues 97-140) and major conformational differences in helices α2/α3 (residues 178-208). The α2 helix is shorter than that in ATXN3L, causing the α3 helix to shift. In the available JOSD2 structure, the corresponding region is not resolved. Interestingly, these regions make important contacts with the proximal (S1’) and distal (S1) ubiquitin units, respectively.

**Figure 2.**
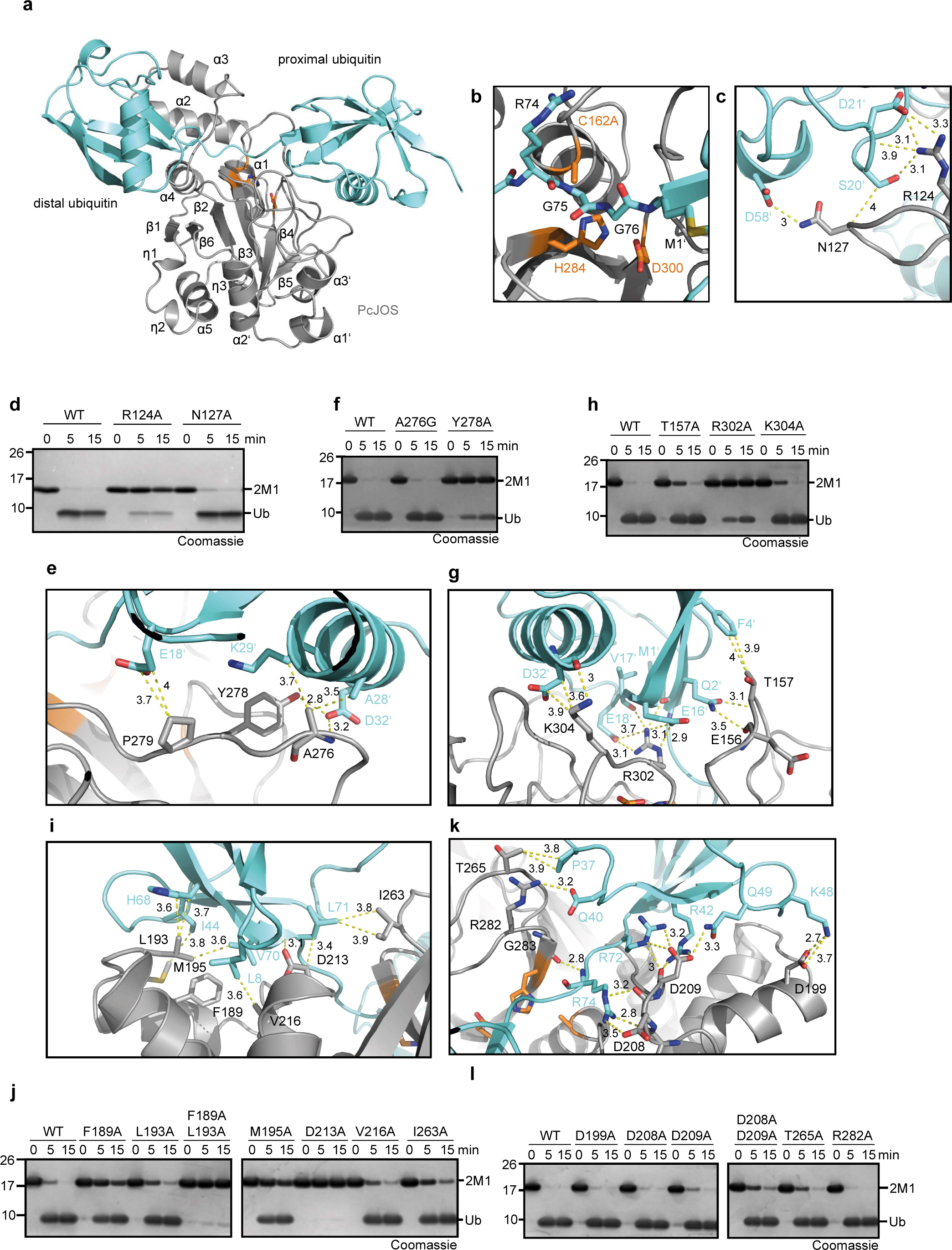
Crystal structure of PcJOS in complex with linear diubiquitin. **a)** Overview of the complex structure in cartoon representation. The catalytic core of PcJOS is shown in grey and ubiquitin in light blue. The catalytic residues are shown as sticks and coloured orange**. b)** Magnification of the PcJOS active site. Important residues are shown as sticks and coloured orange (PcJOS active site) or light blue (ubiquitin C-terminus). The position of ubiquitin C-terminus in relation to the active site residues confirms a cleavage between Arg-74 and Gly-75. **c)** The N-terminal extension is part of the S1’ ubiquitin binding site. Key residues are highlighted as sticks and coloured light blue (ubiquitin) or grey (PcJOS). Hydrogen bonds are indicated by dotted lines. **d)** Activity of PcJOS R124A or N127A against linear linked diubiquitin. **e)** Tyr-278 is a critical part of the S1’ ubiquitin binding site. Tyr-278 and neighbouring residues are highlighted as sticks and coloured light blue (ubiquitin) or grey (PcJOS). Key-interactions are indicated by dotted lines. **f)** Activity of PcJOS A276A or Y278A against linear linked diubiquitin. **g)** Extensive contacts between PcJOS catalytic core and the proximal ubiquitin. Key residues are highlighted as sticks and coloured light blue (ubiquitin) or grey (PcJOS). Interactions are indicated by dotted lines. **h)** Activity of wildtype PcJOS and S1’ site mutants (T157A, R302A and K304A) against linear diubiquitin. **i)** The α2/α3 region is part of the S1 site and recognizes the hydrophobic Ile-44 patch of the distal ubiquitin. Residues involved in these interactions are highlighted as sticks and coloured lightblue (ubiquitin) or grey (PcJOS). Hydrophobic interactions are indicated by dotted lines. **j)** Activity of S1 site mutants shown against linear diubiquitin. **k)** The C-terminus of the distal ubiquitin is stabilized by intensive polar interactions. Residues involved in these interactions are highlighted as sticks and coloured light blue (ubiquitin) or grey (PcJOS). **l)** Mutational analysis of residues stabilizing ubiquitin’s C-terminus.

The S1’ ubiquitin forms an extensive hydrogen-bond network with PcJOS (chain A). Within the N-terminal extension, Arg-124 and Asn-127 form hydrogen bonds with Ser-20’/Asp-21’ and Asp-58’ of the proximal ubiquitin, respectively (Fig 2c). The former contacts are important, since the R124A mutant lost activity almost completely, while N127A did not affect the activity (Fig 2d). To investigate whether the N-terminal region causes M1-linkage specificity by sterically inhibiting other chain types, we tested a truncated version of PcJOS that lacks this region but contains all the structural elements considered important for Josephin catalysis. Unlike the slightly truncated PcJOS^97-304^, which maintained M1-specific cleavage, truncated PcJOS^140-304^ lacking the N-terminal extension was inactive against all tested diubiquitin species and activity-based probes (Supplementary Fig 5f,g). Thus, the M1-specificity of PcJOS is unlikely to be due to the blocking of other chain types, but rather induced by specific linear chain recognition or substrate-assisted catalysis. Additional S1’ contacts are made by loop 275-278, where Tyr-278 contributes a hydrogen bond to Asp-32’, while Ala-276 shows hydrophobic interactions to Ala-28’ and Lys-29’ of the proximal ubiquitin (Fig 2e). As shown in Fig 2f, the Y278A mutant was nearly inactive, while the A28G mutation did not affect activity, suggesting that the Tyr-278 contact is more important for S1’ recognition. In addition, several residues outside this loop contact the proximal ubiquitin (Fig 2g). Arg-302 shows extensive hydrogen bonding with the Glu-16’ and Glu-18’ sidechains and with main chain atoms of Val-17’ and Met-1’. Lys-304 forms a hydrogen bond with the ubiquitin main chain at Asp-32’, and Thr-157 is packed against Phe-4’ of the proximal ubiquitin. Among these contacts, Arg-302 appears to be most important, since the R302A mutant was almost inactive, whereas the K304A and T157A mutants reduced activity only marginally (Fig 2h). To assess whether the side chain contacts of Arg-203 to Glu-16’ and Glu-18’ are both important, wildtype PcJOS was tested against diubiquitin carrying the point mutants E16’A or E18’A. As shown in Supplementary Fig 5h, neither mutant was cleaved, indicating that both hydrogen bonds are required. Overall, the proximal ubiquitin appears to be positioned in a cleavable conformation by a multitude of distinct interactions.

Interactions at the S1 position include recognition of the hydrophobic Ile44-patch by the α2/α3 region and several polar interactions at the C-terminus of ubiquitin close to the scissile bond. Ubiquitin Ile-44 itself contacts Met-195 of PcJOS, whereas the surrounding residues of the patch (Val-70, His-68 and Leu-8) show hydrophobic interactions with Leu-193, Val-216, and Phe-189 of PcJOS (Fig. 2i). Abrogating these interactions individually by the mutations M195A, L193A, V216A, or F189A modestly reduced activity, while the double mutant F189A/L193A was completely inactive (Fig 2j). An additional hydrogen bond between Asp-213 and the main chain of Leu-71 also appeared to be crucial, since the D213A mutant was inactive (Fig 2i,j). Most deubiquitinases recognize and position the cleavable ubiquitin C-terminus through salt bridges and/or hydrogen bonds between Arg-72 and Arg-74 of ubiquitin and acidic or otherwise polar residues of the enzyme. In the PcJOS structure, Arg-72 and Arg-74 contact Asp-209 and Asp-208, respectively (Fig 2k). However, mutations in these residues (D208A, D209A, and D208A/D209A) reduced diubiquitin cleavage only marginally (Fig 2l). In contrast to other DUBs, Arg-72 and Arg-74 not only bind to the side chain carboxyl groups of Asp-208/Asp-209 but also to their main chain carbonyls, which are not affected by mutagenesis. This explanation is supported by experiments using R72A and R74A mutated ubiquitin substrates, which are mostly inert to PcJOS cleavage, demonstrating the importance of these two arginine residues for activity (Supplementary Fig 5h). Additional interactions were observed between Arg-282 and ubiquitin Gln-40, and between Thr-265 and ubiquitin Pro-37, but do not appear crucial, since the mutants T265A and R282A were nearly as active as wildtype PcJOS. In DUBs with conventional cleavage mode, the catalytic histidine residue is usually followed by an aromatic ‘gatekeeper’ residue, which interacts with Gly-75 and – amongst other functions – restricts active site access of substrates without small amino acids at this position ^17^. In UCCs, which cleave after Arg-74, such a gatekeeper role should not be required. Nevertheless, many clippases identified in this study do conserve the aromatic residue (Supplementary Fig 3). In PcJOS, the catalytic His-284 is followed by Phe-285, which contacts Leu-73 of ubiquitin, the residue preceding the cleavage position after Arg-74 (Supplementary Fig 5i). To investigate the importance of this interaction, we tested mutations of the ‘gatekeeper-residue’ and its interaction partner. As shown in Supplementary Fig 5h,j, the activity of the F285A mutant was abolished, and the L73A-mutated ubiquitin was completely resistant to cleavage.

### Structural determinants for clippase-type cleavage

The bacterial Josephin family contains both clippases and conventional DUBs, which raises the question of how the cleavage position is determined. Of particular interest are the family members BpJOS from *Burkholderia pyrrocinia* and PaJOS from *Pigmentiphaga aceris*, since they are closely related, yet differ in their cleavage modes. For conventionally cleaving PaJOS, the catalytic fragment PaJOS^2-265^ was crystallized in a covalent complex with Ub-PA. The structure was solved at a resolution of 1.89 Å with an asymmetric unit containing twelve PaJOS/Ub complexes, which were nearly identical with RMSDs ranging from 0.1 to 0.3 Å. Overall, the structure revealed a Josephin-like papain fold (α1-β6) with an N-terminal extension of a single helix α1’ (Fig 3a). The active site residues were determined as Cys-66, His-187, and Asp-203, confirming their sequence-based prediction. Mutating any of the active site residues to alanine caused a loss of cleavage activity. (Supplementary Fig 6a,b). For the promiscuous clippase BpJOS, the catalytically inactivated full-length protein BpJOS^C69A^ was crystallized in complex with linear diubiquitin. The structure was solved at a resolution of 2.56 Å; The asymmetric unit contained two BpJOS/ubiquitin complexes, each of which showed only one ubiquitin moiety at the S1 position (Supplementary Fig 7a). The electron density suggested that the visible S1-bound ubiquitin corresponds to the C-terminal unit of the diubiquitin substrate, while the N-terminal unit was disordered due to the lack of a defined interface. In both BpJOS chains, residues 15-218 were resolved; their conformation was nearly identical with an RMSD of 0.2 Å over 1308 atoms (Supplementary Fig 7b). The resolved region corresponds to the minimal active fragment of BpJOS, since further truncation of the N-terminal α-helix and the first two β-strands, which are not conserved in PaJOS, caused a complete loss of activity (Supplementary Fig 7c). Overall, the BpJOS structure revealed a Josephin-like papain fold with a core (α1-β5) similar to PaJOS, preceded by a unique N-terminus consisting of one helix and two β-strands (α1’-β2’) (Fig 3b). The active site residues were identified as Cys-96 (Ala-96 in the structure), His-166, and Asp-182; their individual replacement by alanine universally abolished activity (Supplementary Fig 7d,e).

**Figure 3.**
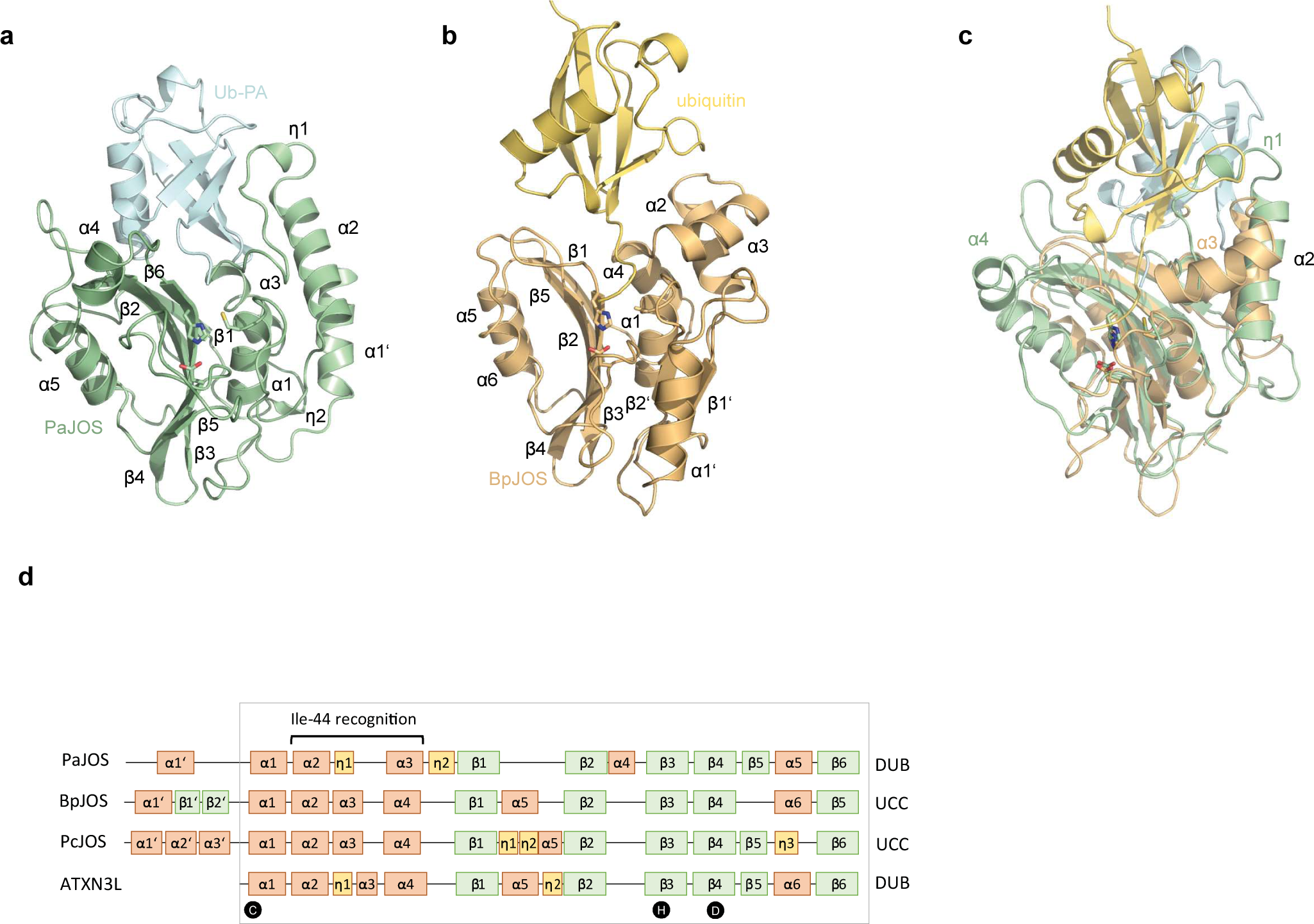
Crystal structures of PaJOS and BpJOS in complex with ubiquitin. **a)** Overview of the PaJOS/Ub-PA complex structure in cartoon representation. The catalytic core of PaJOS is shown in green and ubiquitin in light blue. The catalytic residues are shown as sticks. **b)** Overview of the BpJOS/ubiquitin complex structure in cartoon representation. The catalytic core of BpJOS is shown in orange and ubiquitin in yellow. The catalytic residues are shown as sticks. **c)** Structural superposition of the DUB PaJOS and the UCC BpJOS structures shown in a/b). The superposition is based on the catalytic domains which align with a RMSD of 1.29 Å over 522 atoms. Secondary structure elements differing in the structures are numbered. **d)** Schematic overview over secondary structure elements of bacterial Josephins in comparison to human ATXN3L. The catalytic core domain is indicated by a grey box. The position of the catalytic residues is marked by black circles.

In accordance with their sequence similarity, the catalytic domains of PaJOS and BpJOS can be superimposed with an RMSD of 1.29 Å over 522 atoms. (Fig 3c). However, the bound S1 ubiquitin molecules do not superimpose, since they are bound in a different orientation, which ultimately causes displacement of their C-termini and thus a shifted cleavage position. The differences in ubiquitin binding appear to be caused by subtle changes in protease structure. Both enzymes bind the Ile-44 patch of S1-ubiquitin through homologous regions corresponding to α2/η1/α3 in PaJOS and α2/α3/α4 in BpJOS (Fig 3d), with poorly conserved sequences and major structural differences. In BpJOS, this region forms a rigid helical structure and contacts the Ile-44 patch via Val-96, Leu-98, and Phe-116 (Fig 4a). Here, the α3 helix provides rigidity, but does not directly contact ubiquitin. By contrast, the corresponding region of PaJOS is more flexible; α2 is extended by the short helix η1 and connected via an unstructured loop to α3. The ubiquitin Ile-44 patch is contacted by Phe-96, Ile-97, and Leu-104 (Fig 4b). Mutagenesis of these hydrophobic residues showed that for both enzymes, ubiquitin cleavage is highly dependent on these contacts (Fig 4c,d). The structural differences in the Ile-44 recognition regions are likely to cause the different orientation of the bound ubiquitin. Among other steric problems, the S1 ubiquitin bound by BpJOS would clash with η1 of PAC.

**Figure 4.**
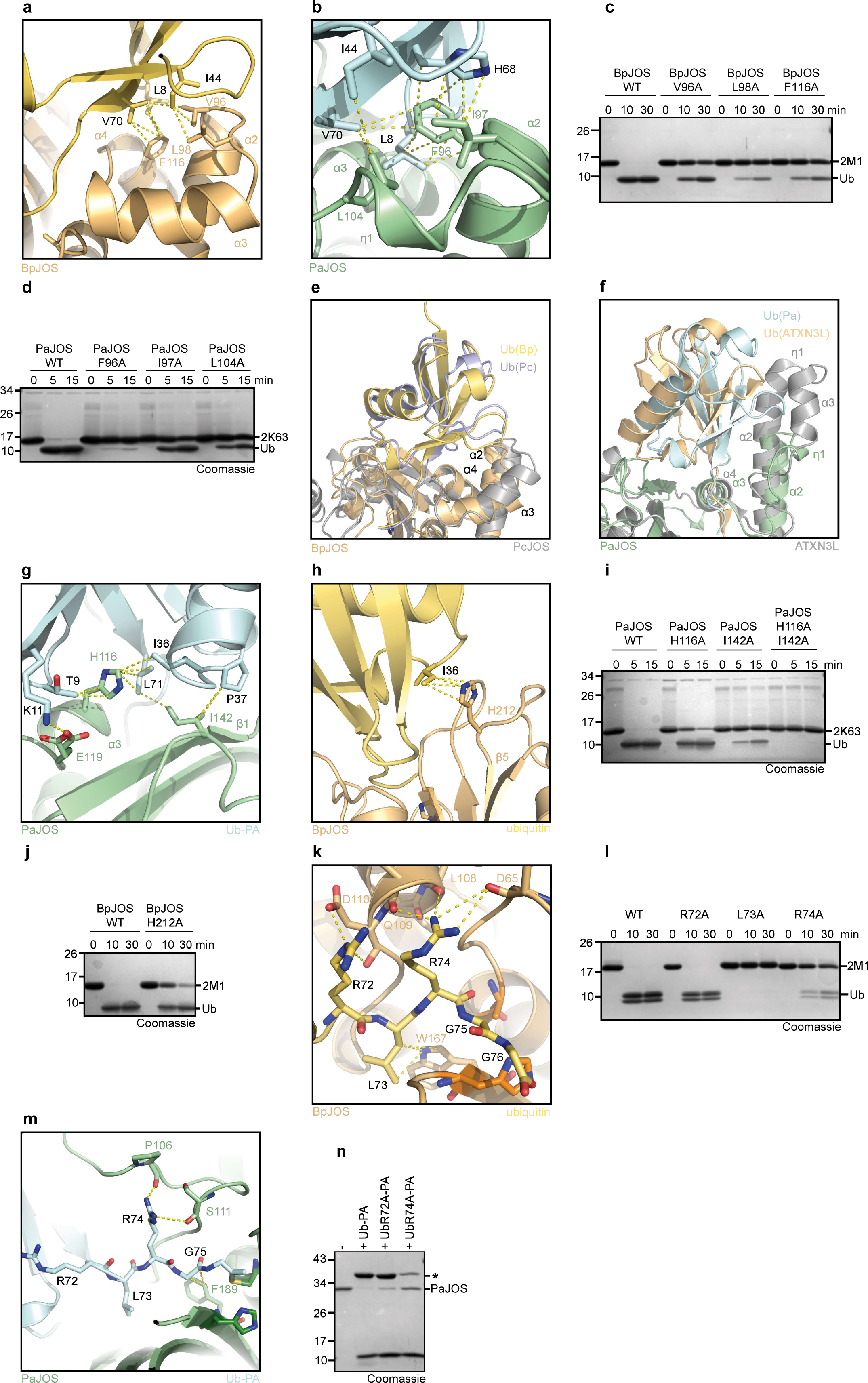
Differential S1 ubiquitin binding by Josephin family DUBs and UCCs. **a)** Recognition of ubiquitin’s Ile-44 patch by the α2/α3/α4 region of BpJOS. Residues involved in these interactions are highlighted as sticks and coloured yellow (ubiquitin) or light orange (BpJOS). Hydrophobic interactions are indicated by dotted lines. **b)** Recognition of ubiquitin’s Ile-44 patch by the α2/η1/α3 region of PaJOS. Residues involved in these interactions are highlighted as sticks and coloured light blue (ubiquitin) or green (PaJOS). Hydrophobic interactions are indicated by dotted lines. **c** Activity of wildtype BpJOS or S1 site mutants against linear diubiquitin. **d)** Activity of wildtype PaJOS or S1 site mutants against K63-linked diubiquitin. **e)** Structural superposition of the UCCs BpJOS (orange) and PcJOS (grey) structures. The superposition is based on the catalytic domains which align with a RMSD of 2.362 over 619 atoms. The bound ubiquitin molecules are in an almost identical orientation and the conserved secondary structure elements involved in ubiquitin binding are numbered. **f)** Structural superposition of the DUBs PaJOS (green) and ATXN3L (grey, PDB: 3O65) structures. The superposition is based on the catalytic domains which align with a RMSD of 2.37 over 520 atoms. The bound ubiquitin molecules are in a similar orientation and the secondary structure elements involved in ubiquitin binding are numbered. **g,h)** Ile36-patch recognition by PaJOS (g) or BpJOS (h). Residues involved in the interaction are highlighted as sticks. **i)** Activity of wildtype PaJOS or S1 site mutants against K63-linked diubiquitin. **i)** Activity of wildtype BpJOS or H212A against linear diubiquitin. **k)** Recognition of ubiquitin’s C-terminus by BpJOS. Residues involved in these interactions are highlighted as sticks and coloured yellow (ubiquitin) or light orange (BpJOS). Important interactions are indicated by dotted lines. **l)** Activity of BpJOS against ubiquitin mutants. N-terminally His-tagged and mutated linear diubiquitin was incubated with BpJOS for the indicated timepoints. **m)** Recognition of ubiquitin’s C-terminus by PaJOS. Residues involved in these interactions are highlighted as sticks and coloured lightblue (ubiquitin) or green (PaJOS). Important interactions are indicated by dotted lines. **n)** Activity-based probe reaction of PaJOS with wildtype, R72A or R74A Ub^1-75^-PA. The reaction was stopped after 3h.

To investigate if the two ubiquitin binding modes are conserved in other Josephin UCCs and DUBs, we compared the two available clippase structures BpJOS and PcJOS, whose catalytic domains can be superimposed with an RMSD of 2.36A over 619 atoms. As shown in Fig 4e, the position of the α3 helices of BpJOS and PcJOS are conserved and the orientation of the bound ubiquitin is nearly identical (Fig 4e). Conversely, both the conventionally cleaving PaJOS and the eukaryotic DUB ATXN3L use their α2 helices, each of them extended by a short η1 helix, to position the S1 ubiquitin ready for DUB-cleavage (Fig 4f). Taken together, the available data suggest that Josephin-type UCCs use the helices following the catalytic cysteine in a conserved way to position the S1 ubiquitin for clippase cleavage, while bacterial Josephin-type DUBs resemble their eukaryotic counterparts. Both PaJOS and BpJOS have a second ubiquitin-binding interface contacting the Ile-36 patch. In PaJOS, residues His-116 and Ile-142 engage in hydrophobic interactions with Thr-9, Ile-36, Pro-37, and Leu-71 of the S1 ubiquitin (Fig 4g). In BpJOS, His-212 interacts with Ile-36 of ubiquitin (Fig 4h). In both cases, these interactions are important for full activity, as demonstrated by mutational analysis (Fig 4i,j). Owing to the different ubiquitin orientations, Ile-36 recognition involves different regions of PaJOS and BpJOS (Supplementary Fig 6c). While the DUB-typical Ile-36 recognition is conserved between PaJOS and ATXN3L (Supplementary Fig 6d), the recognition in UCC-orientation differs between the two available clippase structures. The loop, which in BpJOS contacts Ile-36, is shorter in PcJOS. Instead, Ile-263 from a neighboring loop contacts Leu-71 in ubiquitin. Despite their different sequence positions, the contact residues occupy a similar space, resulting in a conserved orientation of the bound ubiquitin (Supplementary Fig 7f).

Since UCCs bind the ubiquitin C-terminus shifted by two residues relative to DUBs, major differences in positioning the ubiquitin tail are expected. In PcJOS, Arg-72 and Arg-74 are crucially stabilized by hydrogen bonds with backbone atoms (Fig 2k, Supplementary Fig 5h); an analogous binding mode was observed for BpJOS. Arg-74 of ubiquitin, the residue directly preceding the scissile bond, forms strong hydrogen bonds with backbone atoms of Asp-65, Leu-108, and Gln-109 of BpJOS. Another hydrogen bond was observed between Arg-72 of ubiquitin and Asp-110 of BpJOS (Fig 4k). Accordingly, linear R74A diubiquitin was hardly cleaved by BpJOS, while the R72A substitution had no visible effect on cleavage (Fig 4l). Leu-73, the hydrophobic residue between Arg-72 and Arg-74, contacts Trp-167 of BpJOS (and Phe-285 of PcJOS) and is crucial for catalysis (Fig 4k,l). In the conventionally cleaving and K63-prefering PaJOS, Arg-74 of ubiquitin is also stabilized by a main chain hydrogen bond, albeit to a different region of the protease (Pro-106 and Ser-111). Arg-72 does not show strong interactions (Fig 4m). Accordingly, PaJOS reacted with R72A-PA, whereas its reaction with R74A-PA was strongly impaired (Fig 4n). Unlike BpJOS, PaJOS does not interact with Leu-73, which appears to be a clippase-specific requirement. Instead, the aromatic gatekeeper residue Phe-188 stabilizes Gly-75, as is commonly observed in conventional DUBs (Fig 4m). The effect of the F188A mutant on diubiquitin cleavage was modest (Supplementary Fig 6e). The shifted ubiquitin C-terminus is shown in Supplementary Fig 7g, which highlights that although both require stabilization of R74, the interacting residues are located differently.

### Predictability of the cleavage position

Among the bacterial Josephins described so far are four linkage-promiscuous UCCs, one M1-specific UCC, and two conventionally cleaving DUBs. To expand our knowledge of clippases, search for additional linkage-specificities, and test whether structural insights into ubiquitin positioning are predictive of the cleavage mode, we performed a more comprehensive bioinformatical search for additional candidates. We selected a set of eight representative proteins for experimental validation while avoiding closely related sequences (PtJOS, KrJOS, MtJOS, MxJOS, ScJOS, MlJOS, Pc2JOS, and CsJOS; see Supplementary Table 1 for accession data and Supplementary Fig 3 for alignment). By subjecting the catalytic domains of these proteins to Alphafold modelling in complex with ubiquitin, seven plausible models with bound ubiquitin were obtained (Supplementary data file 1). KrJOS yielded an unsatisfactory model with a poor complex confidence score (iptm+ptm < 0.35). A superposition of the resulting models with the ubiquitin-bound structures of a clippase (BpJOS) and a DUB (PaJOS) revealed that only MxJOS had ubiquitin modelled in the DUB-typical arrangement, whereas the other six models showed a clippase-oriented S1 ubiquitin (Fig 5a, Supplementary Fig 8).

**Figure 5.**
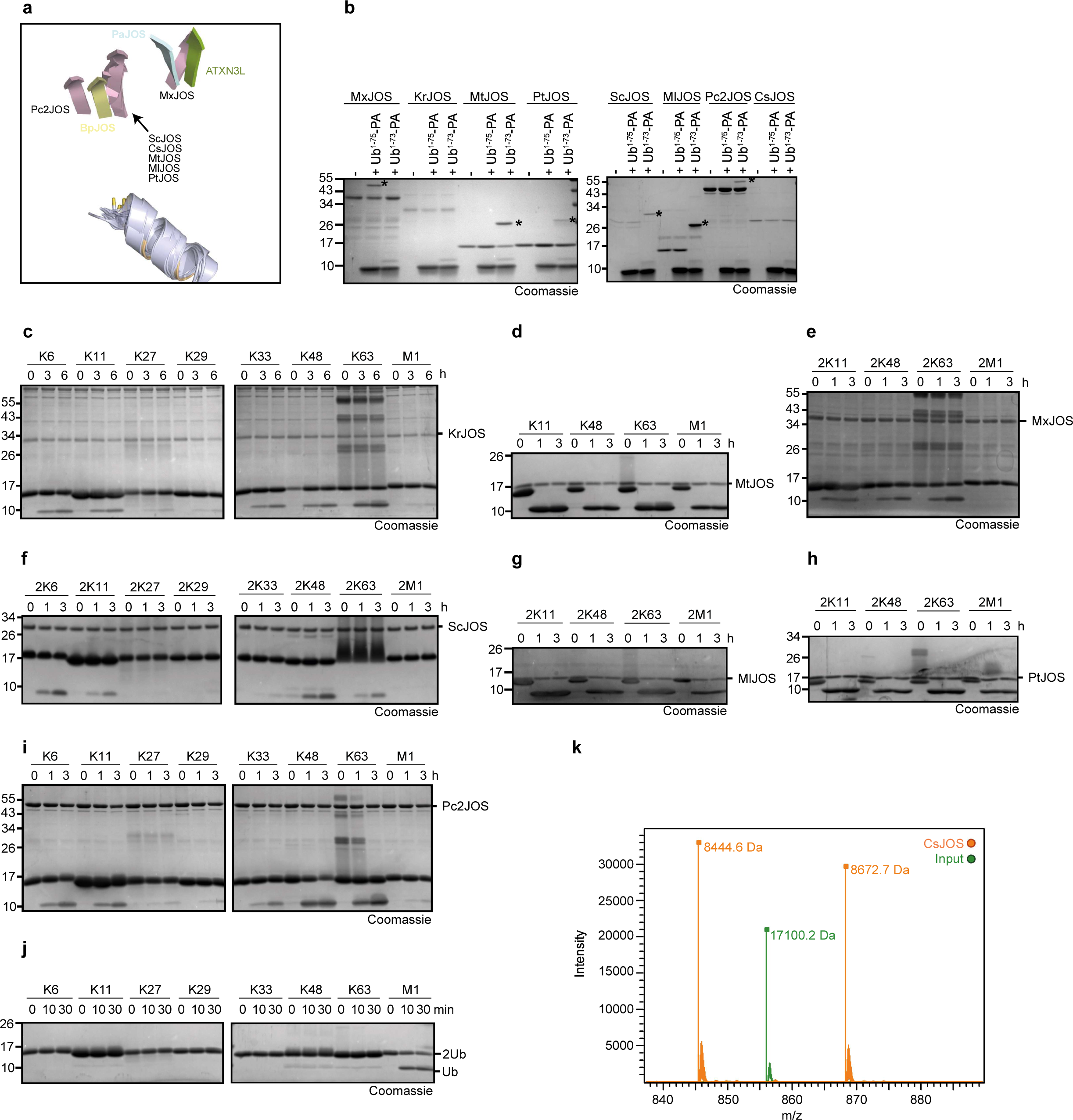
Predictability of UCC and DUB cleavage. **a)** Structures of divergent bacterical Josephins were predicted using AlphaFold2 in complex with ubiquitin. The predicted structures and the crystal structures of BpJOS and ATXN3L (PDB: 3O65) were superimposed onto PaJOS. For each model, only the catalytic cysteine-containing helix and β-3 of the bound ubiquitin are shown. Comparison of the relative orientation of the bound ubiquitin β-strand (pink) to PaJOS (light blue)/ATXN3L (green) or BpJOS (yellow) allows to predict for DUB or clippase cleaving activity. **b)** Determination of the cleavage position by activity-based probes. The new candidates shown in a) were incubated with Ub^1-75^ or Ub^1-73^-PA probes for 18h. Asterisks (*) mark the shifted bands after the reaction. **c-j)** Linkage specificity analysis of bacterial Josephin-DUBs. A panel of diubiquitin chains was treated with 10 µM KrJOS (c), 5µM MtJOS (d), 5 µM MxJOS (e), 5 µM ScJOS (f), 5 µM MlJOS (g), 5 µM PtJOS (h), 5 µM Pc2JOS (i) or 50 nM CsJOS (j) for the indicated time points. **k)** Intact mass spectrometry of M1-linked diubiquitin cleaved by CsJOS. The m/z ratio is depicted on the x-axis, while the deconvoluted masses are shown next to the respective peaks. The input sample shows a single 17100 Da peak (green) corresponding to the monoisotopic mass of diubiquitin, which gets cleaved by CsJOS to two different products (blue) corresponding to the mono-isotopic mass of ubiquitin ± a GlyGly peptide (8672 / 8444 Da).

For all eight candidates, catalytic domains were expressed in *E. coli*, purified, and tested for their cleavage mode and possible linkage specificity. KrJos, the candidate without a convincing Alphafold model, was poorly expressed and did not react with the UCC-specific probe Ub^1-73^-PA or with the DUB-specific Ub^1-75^-PA (Fig 5b). For all candidates with high-confidence complex models, the experimentally determined cleavage modes matched the structure-based predictions: MxJOS reacted only with conventional Ub^1-75^-PA, whereas MtJOS, PtJOS, ScJOS, MlJOS, and Pc2JOS reacted only with the clippase probe Ub^1-73^-PA (Fig 5b). All these enzymes cleaved diubiquitin species of various linkage types without much selectivity (Fig 5c-i). In contrast, CsJOS from *Chlamydiales bacterium ST3* did not react with any of the probes and exclusively cleaved linear diubiquitin (Fig 5b,j). Since linkage-specific DUBs often do not react with probes, we analyzed the M1-linked diubiquitin digestion by intact mass spectrometry and found that CsJOS cleaves in clippase mode, as predicted from the structural model (Fig 5k).

### Clippase toxicity

Bacterial Josephins with clippase activity are found in diverse bacterial phyla, including Pseudomonadota, Chlamydiota, Myxococcota, Acidobacteriota, and Cyanobacteria (Supplementary Table 1), but their occurrence within these phyla is restricted to a few species, excluding the major human pathogens. To investigate if the destructive deubiquitination performed by UCCs is more damaging to host cells than conventional deubiquitination, we expressed similar levels of the promiscuous clippase BpJOS, the M1-specific clippase PcJOS, and the highly-active promiscuous deubiquitinase USP2 in human HEK293 cells (Fig 6a). While BpJOS shows significant toxicity starting 15h post transfection, which increases over the next 30h, the conventional DUB USP2 and the M1-specific clippase PcJOS showed modest toxicity starting 30h post transfection. Simultaneous treatment with the caspase-inhibitor zVAD-fmk had no effect on DUB-induced cell death (Fig 6b). The anti-Flag Western blot shows the PcJOS and USP2 levels to be as least as high as the BpJOS level (Fig 6c).

**Figure 6.**
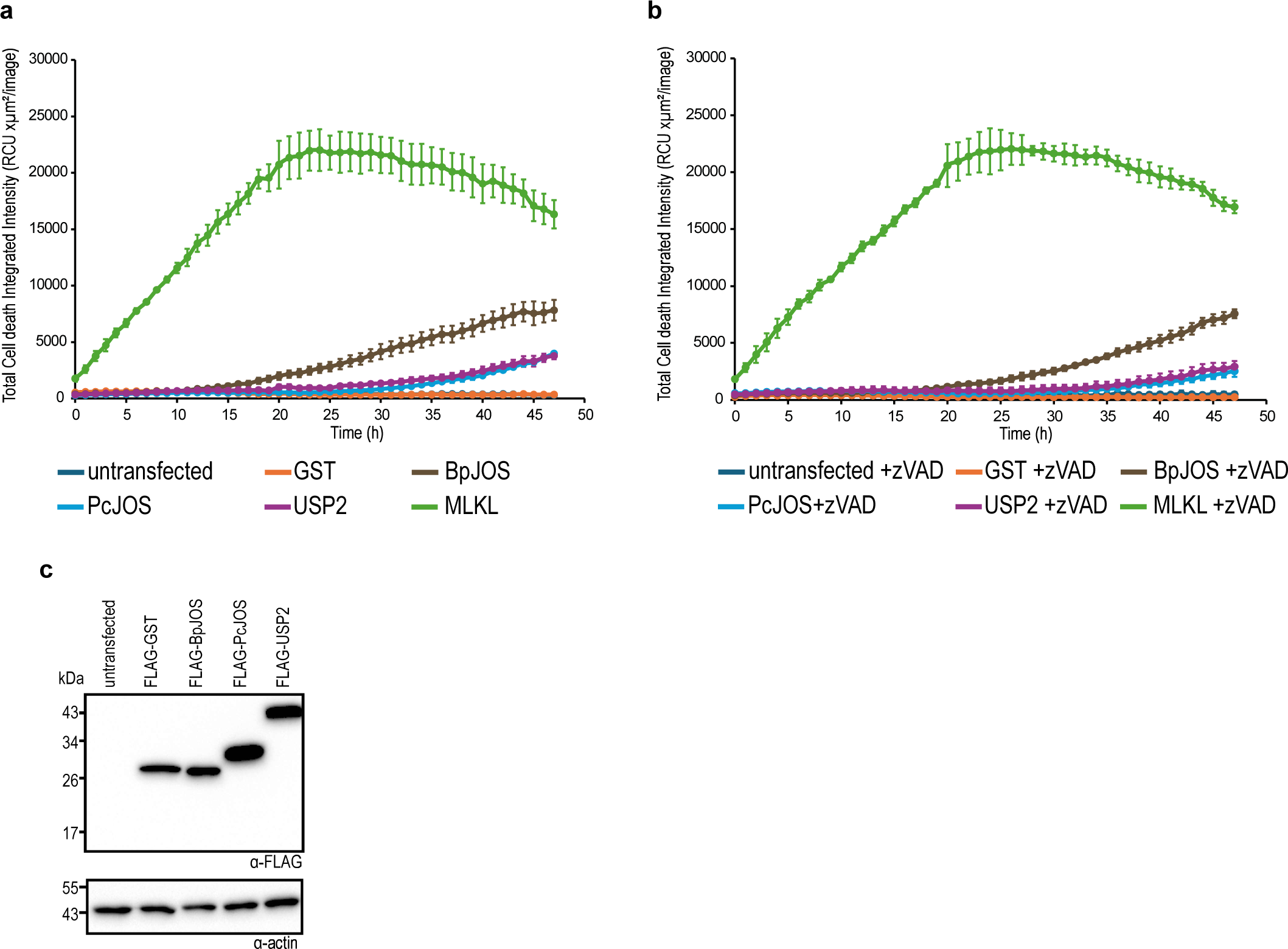
Clippase toxicity. **a,b**) Lytic cell death in the absence (a) or presence (b) of zVAD-FMK, observed by live-cell imaging using DRAQ-7 incorporation as a metric for lytic cell death. Time course data are shown for BpJOS (brown), PcJOS (light blue), USP2 (purple), GST (orange), untransfected cells (dark blue) and cells transfected with the necroptosis inducer MLKL-NT (green) as a positive control. Data points represent the mean of four technical replicates. Error bars correspond to the standard deviation. **c)** Expression control by anti-FLAG Western blot, showing similar protein amounts for the different deubiquitinases.

## Discussion

Intracellular bacteria use a variety of effectors to counteract host ubiquitin-based defense mechanisms. Besides deubiquitinases, which can reverse the action of host E3 ligases^11^, effector types have been described that can prevent access of ligases to bacteria^9^, methylate bacterial surface lysines^28^, impair host ubiquitin through deamidation^10^, or destroy host E2 enzymes by crosslinking them to ubiquitin^29^. Ubiquitin C-terminal clippases (UCCs) are new weapons in the bacterial armory, which combine the deubiquitination of undesirable targets with the irreversible destruction of the modifier and the protection of the modification site against re-modification. At first glance, such effectors appear ideal for bacterial subversion of the host ubiquitin-based defense, which raises the question of why not more bacteria use clippases than conventional DUBs. One possible explanation is that many bacteria rely on ubiquitin ligase activity, which makes the wholesale destruction of ubiquitin disadvantageous. Moreover, high clippase activity maintained over a longer period causes cell death (Fig 6), possibly due to the depletion of functional ubiquitin and/or the clogging of ubiquitin recognition components by dysfunctional truncated ubiquitin. Among the clippase-encoding bacteria, only *Simkania negevensis* has been shown to infect human cells^30^, but the expression levels of SnJOS1 and SnJOS2 during infection are very low^18^ and both enzymes have rather modest activities (Fig 1b). At the other end of the spectrum is the highly active BpJOS from *Burkholderia pyrrocinia*, a bacterium that is occasionally found in cystic fibrosis patients but lacks an infection model^31^.

Bacterial Josephin-family clippases are the first naturally occurring enzymes with ubiquitin-directed clipping (UCC) activity. The leader peptidase Lb^pro^ of foot- and-mouth disease virus (FMDV) cleaves, besides the viral polyprotein, the ubiquitin-like modifier ISG15 before the terminal GlyGly ^19^. Based on Lb^pro^, a variant enzyme Lb^pro^* has been engineered to show reduced modifier specificity, including ubiquitin and NEDD8 ^20^. However, its UCC activity remains several orders of magnitude lower than that of BpJOS (Supplementary Fig 4c). Unlike the singleton Lb^pro^, bacterial clippases belong to an extended family containing both UCC and DUB members. These favorable circumstances allowed us to address the structural changes required to shift the cleavage site by two positions. At least within the Josephin family, the crucial factor is the orientation of the bound S1 ubiquitin, whose Ile-44 patch is recognized by the variable α2/α3/α4 region located between the helix (α1) and the first β-strand of the papain fold (Fig 3). The available UCC and DUB structures differ markedly within this region, resulting in clearly distinct ubiquitin orientations. Interestingly, the α2/α3/α4 region shows considerable variability among different clippases, both in sequence and (predicted) structure, while the orientation of the bound S1 ubiquitin is predicted to be better conserved (Fig 5a) and was found to be highly predictive of the experimentally determined cleavage mode (Fig 5b,k).

Besides their insights into DUB evolution and bacterial defense mechanisms, the Josephin-type clippases also offer new possibilities for experimentally studying the ubiquitin system in general. Engineered multi-UBL clippase Lb^pro^* has already been successfully applied to study branched ubiquitin chains ^20^. After treatment with clippases, each ubiquitination site leaves a diGly remnant on a lysine residue, which can be identified by mass spectrometry. In the case of ubiquitin, the detection of multiple remnants on a single ubiquitin unit proves the presence of branches and allows to quantify them^20^. However, Lb^pro^* does not discriminate between ubiquitination, NEDDylation, and ISGylation, as all of these modifications are cleaved equally well. Moreover, being derived from a physiological deISGylase, Lb^pro^* has no specificity for ubiquitin linkage types, and thus does not yield information on the nature of the chain branches. Bacterial UCCs may be key to addressing these limitations. BpJOS is a highly active pan-linkage UCC that completely spares ISGylation sites (Supplementary Fig 4), allowing discrimination between ubiquitin- and ISG15-derived diGly remnants, which co-occur under stimulation of immunity. Since BpJOS also cleaves NEDD8-modifications in clippase mode, it cannot be used in its present form to discriminate between ubiquitination and NEDDylation sites. However, the structural data provided here should allow the engineering of modifier-specific clippases. Another benefit of the bacterial Josephin-type clippases is the presence of linkage-specific enzymes. Among the eleven UCCs characterized in this study, five are highly active linkage-promiscuous enzymes, two are strictly M1-specific, and the rest exhibits modest linkage preferences without real specificity (Figs 1,5 and Ref ^18^). Current protein databases contain several dozen additional UCC candidates, and new bacterial sequences are being added continuously. The available sequence and structural diversity will be instrumental in the identification and engineering of additional clippases with new specificities.

## Materials & Methods

### Sequence Analysis

Sequence alignments were generated using the MAFFT package^32^. Generalized profiles were derived from multiple alignments using pftools3 ^26^ and searched against the Uniprot database (https://www.uniprot.org) and the NCBI microbial genome reference sequence database (https://www.ncbi.nlm.nih.gov/genome/microbes) using pfsearchV3 ^33^. HMM-to-HMM searches were carried out using the HHSEARCH method ^34^. Structure predictions were run using a local installation of Alphafold 2.3 ^35^. For structure comparisons, the DALI software was used ^27^.

### Cloning & Mutagenesis

All coding regions of bacterial Josephin DUBs, except SnJOS2, were obtained by gene synthesis (IDT) and cloned into pOPIN-S vector ^36^ using the In-Fusion HD Cloning Kit (Takara Clontech). SnJOS2 was amplified from *S. negevensis* genomic DNA (kind gift of Vera Kozjak-Pavlovic, Julius Maximilian University, Würzburg) and cloned accordingly. A codon optimized diubiquitin was obtained by gene synthesis and cloned into pOPIN-B vector ^36^. Lbpro* was obtained by gene synthesis and cloned into pOPIN-K vector ^36^. Point mutations were introduced using the QuikChange Lightning kit (Agilent Technologies). Constructs for ubiquitin-PA purification (pTXB1-ubiquitin^1–75^) and USP21 were kind gifts of David Komander (WEHI, Melbourne). USP2 was a kind gift of Malte Gersch (Technical University of Dortmund). MLKL was a kind gift of Xiaodong Wang (Hebei Agricultural University, Baoding). Coding sequences for BpJOS and PcJOS were optimized for human codons and obtained by gene synthesis (IDT). For the expression in human cells, GST^59-712^, BpJOS, PcJOS^70-324^ and USP2^258-605^ were cloned with an N-terminal FLAG (DYKDDDDK) tag and MLKL^1-180^ with a C-terminal FLAG-tag into pcDNA™5/FRT/TO vector (Invitrogen™).

### Protein expression & purification

The bacterial and human Josephins were expressed from the pOPIN-S vector with an N-terminal 6His-SMT3-tag. ISG15^79-165^ and ubiquitin were expressed from the pOPIN-B vector and Lb^pro^* from pOPIN-K vector with an N-terminal 6His-tag or 6His-GST-tag respectively. *Escherichia coli* (Strain: Rosetta (DE3) pLysS) were transformed with the respective constructs and 2-6⍰l cultures were grown in LB medium at 37⍰°C until the OD_600_ of 0.8 was reached. The cultures were cooled down to 18⍰°C and protein expression was induced by addition of 0.1⍰mM isopropyl β-d-1-thiogalactopyranoside (IPTG).

The expression of selenomethionine substituted PcJOS was carried out as described previously ^37^: In brief, the expression cultures were grown in M9 minimal medium supplemented with thiamine vitamin (0.0001% w/v final concentration) until the OD600 of 0.8 was reached. The cultures were mixed with feed-back inhibition amino acid mix (0.5⍰g/l leucine, isoleucine, valine, selenomethionine and 1 g/l lysine, threonine, phenylalanine), induced with 0.2⍰mM IPTG and cooled down to 18⍰°C. After 16⍰h, the cultures were harvested by centrifugation at 5000⍰×⍰*g* for 15⍰min.

After 16⍰h, the cultures were harvested by centrifugation at 5000⍰×⍰*g* for 15⍰min. After freeze thaw, the pellets were resuspended in binding buffer (300⍰mM NaCl, 20⍰mM TRIS pH 7.5, 20⍰mM imidazole, 2⍰mM β-mercaptoethanol) containing DNase and lysozyme, and lysed by sonication using 10 s pulses with 50 W for a total time of 10 min. Lysates were clarified by centrifugation at 50,000⍰×⍰g for 1⍰h at 4⍰°C and the supernatant was used for affinity purification on HisTrap FF columns (GE Healthcare) according to the manufacturer’s instructions. With the exception of ubiquitin and MxJOS all 6His-Smt3, 6His-GSTand 6His-tags were removed by incubation with SENP1^415-644^ or 3C protease respectively. The proteins were simultaneously dialyzed in binding buffer. The liberated affinity-tag and the His-tagged SENP1 were removed by a second round of affinity purification with HisTrap FF columns (GE Healthcare). All proteins were purified with a final size exclusion chromatography (HiLoad 16/600 Superdex 75pg) in 20⍰mM TRIS pH 7.5, 150⍰mM NaCl, 2⍰mM dithiothreitol (DTT), concentrated using VIVASPIN 20 Columns (Sartorius), flash frozen in liquid nitrogen, and stored at −80⍰°C. Protein concentrations were determined using the absorption at 280 nm (A_280_) using the proteins’ extinction coefficients derived from their sequences.

### Enzymatic generation of activity-based probes

Wildtype Ub^1-75^-PA, the R72A / R74A mutants and ISG15^79-154^-PA were expressed as C-terminal intein fusion proteins. The intein fusion proteins were affinity purified in buffer A (20⍰mM HEPES, 50⍰mM sodium acetate pH 6.5, 75⍰mM NaCl) from clarified lysates using Chitin Resin (New England Biolabs) following the manufacturer’s protocol. On-bead cleavage was performed by incubation with cleavage buffer (buffer A containing 100⍰mM MesNa (sodium 2-mercaptoethanesulfonate)) for 24⍰h at room temperature (RT). The resin was washed extensively with buffer A and the pooled fractions were concentrated and subjected to size exclusion chromatography (HiLoad 16/600 Superdex 75pg) with buffer A. To synthesize the propargylated probe, 300⍰µM Ub/Ubl-MesNa were reacted with 600⍰mM propargylamine hydrochloride (Sigma Aldrich) in buffer A containing 150⍰mM NaOH for 3⍰h at RT. Unreacted propargylamine was removed by size exclusion chromatography and the probes were concentrated using VIVASPIN 20 Columns (3⍰kDa cutoff, Sartorius), flash frozen and stored at −80⍰°C.

### Chemical synthesis of activity-based probes

Ub^1-73^ and Nedd8^1-73^ were synthesized on a Syro II MultiSyntech Automated Peptide synthesizer using standard 9-fluorenylmethoxycarbonyl (Fmoc) based solid phase peptide chemistry on a 20 µmol scale as described previously ^38^. Here, the N-terminal methionine (and position 50 methionine in Nedd8) was replaced by the known isostere norleucine. After N-terminal Boc protection, sidechain-protected Ub or Nedd8 was released from the resin with HFIP/DCM (2.5 ml, 1/4, v/v, 3x 20 min). 10 µmol of the cleaved peptide was dissolved in TFE/CHCl (6 ml, 1/1, v/v) and cooled to -10°C to prevent racemization ^39^. PA (6.40 µl, 100 µmol, 5 eq.), EDC.HCl (19.2 mg, 100 µmol, 5 eq.), and HOBt (15.3 mg, 100 µmol, 5 eq.) were added and stirred for 10 min at -10°C, then overnight at RT. After confirming full conversion, the excess solvent was evaporated and global deprotection was performed with TFA/Tis/H_2_O/Phenol (6 ml, 90/2.5/5/2.5, v/v) for 3 hours. The mixture was added to cold ether/pentane (40 ml, 1/3, v/v), precipitating the product, which was isolated by centrifugation (3500 rpm, 5 min, 4°C) and washed with cold diethyl ether (3x). The pellet was redissolved in DMSO (1.5 ml) and diluted in water (40 ml) for purification by RP-HPLC. The pure fractions were lyophilized affording the title compound Ub^1-73^-PA and Nedd8^1-73^-PA as white powders. Ub^1-73^-PA (7.15 mg, 0.86 µmol, 8.6%). MS ES+ (amu) calculated: M+H+ = 8314 Da, deconvoluted mass found: M+H+ = 8314 Da. Nedd8^1-73^-PA. MS ES+ (amu) calculated: M+H+ = 8289 Da, deconvoluted mass found: M+H+ = 8289 Da.

### Chain generation

Untagged Met1-linked diubiquitin was expressed as a linear fusion protein and purified by ion exchange chromatography and size exclusion chromatography. Wildtype 6His-tagged Met1-linked diubiquitin and mutants were expressed as linear fusion proteins and purified by HisTrap affinity purification and size exclusion chromatography. K11-, K48-, and K63-linked ubiquitin chains were enzymatically assembled using UBE2SΔC (K11), CDC34 (K48), and Ubc13/UBE2V1 (K63) as previously described^40, 41^. In brief, ubiquitin chains were generated by incubation of 1⍰µM E1, 25⍰µM of the respective E2, and 2⍰mM ubiquitin in reaction buffer (10⍰mM ATP, 40⍰mM TRIS (pH 7.5), 10⍰mM MgCl_2_, 1⍰mM DTT) for 18⍰h at RT. The respective reactions were stopped by 20-fold dilution in 50⍰mM sodium acetate (pH 4.5) and chains of different lengths were separated by cation exchange using a Resource S column (GE Healthcare). Elution of different chain lengths was achieved with a gradient from 0 to 600⍰mM NaCl.

### Intact mass analysis

Samples were analyzed by the Proteomics Facility (CECAD, Cologne) on a Shimadzu Nexera X2 coupled to a TripleTOF 6600 using a DuoSpray ion source heated to 150 °C (both Sciex). Samples were separated on a Jupiter C4 column (150 cm length, 1 mm inner diameter, Phenomenex) using a 5 min isocratic gradient of 20 % acetonitrile with 0.2 % formic acid. After 5 min, washing was performed by increasing acetonitrile concentration to 85 % for 3 min, followed by re-equilibration to starting conditions. Acquisition was performed in positive MS1 between 600 and 1600 m/z with a declustering potential of 10. System control and data acquisition were done using Analyst TF 1.8.1, which was also used to export integrated spectra of relevant peaks. Afterwards, annotation of monoisotopic masses and subsequent deconvolution of charge clusters was done in mMass 5.5 ^42^.

### Crystallization

Catalytic inactive PcJOS (± selenomethionine substitution) and linear linked diubiquitin were mixed in a 1:1.1 ratio and crystallized using sitting drop vapor diffusion with commercially available sparse matrix screens. 96 well crystallization plates containing 30⍰µl of the respective screening conditions were mixed with 10⍰mg/ml protein in the ratios 1:2, 1:1 and 2:1 in 300⍰nl drops. Initial crystals of the native complex appeared in MIDAS G6 (35 % v/v glycerol ethoxylate, 0.2 M lithium citrate) at 20°C and were cryoprotected with Perfluoropolyether. Best diffracting crystals of the selenomethionone substituted complex were harvested from Morpheus B7.

100⍰µM PaJOS were incubated with 200⍰µM ubiquitin-PA for 18⍰h at 4⍰°C. Unreacted PaJOS and Ub-PA were removed by size exclusion chromatography. The covalent PaJOS /Ub-PA complex (10⍰mg/ml) was crystallized using the vapor diffusion with commercially available sparse matrix screens. Crystallization trials were set up with drop ratios of 1:2, 1:1, 2:1 protein solution to precipitant solution with a total volume of 300⍰nl. Initial crystals appeared in Crystal A6 (0.2 M magnesium chloride, 0.1 M TRIS pH 8.5, 30 % w/v PEG4000) at 20°C. These crystals were optimized by gradually changing the pH and PEG4000 concentration using 48-well MRC plates with 80 µl reservoir solutions and 3 µl drops (protein/precipitant ratios: 2:1, 1:1 and 1:2). Best diffracting crystals were harvested from a condition containing 0.2 M magnesium chloride, 0.1 M TRIS pH 9, 30 % w/v PEG4000.

Catalytic inactive BpJOS and linear linked diubiquitin were mixed in a 1:1.1 ratio and crystallized using sitting drop vapor diffusion with commercially available sparse matrix screens. 96 well crystallization plates containing 30⍰µl of the respective screening conditions were mixed with 10⍰mg/ml protein in the ratios 1:2, 1:1 and 2:1 in 300⍰nl drops. Initial crystals of the native complex appeared in Wizard A1 (20 % w/v PEG8000, 0.1 M CHES pH 9.5) at 20°C. These crystals were optimized by gradually changing the pH and PEG8000 concentration using 48-well MRC plates with 80 µl reservoir solutions and 3 µl drops (protein/precipitant ratios: 2:1, 1:1 and 1:2). Best diffracting crystals were harvested from a condition containing 0.1 M CHES pH 9.5; 22 % w/v PEG8000 and were cryoprotected with reservoir solution containing 20 % w/v glycerol

### Data collection, phasing, model building, and refinement

All diffraction data were processed using XDS ^43^. Molecular replacement structure solution was performed using PHASER ^44^. Refinement was achieved by phenix.refine and REFMAC, and model building using the program COOT ^45-47^. The tetragonal crystal form of PcJOS was used for selenomethionine phasing with data collected at beamline X06SA, Swiss Light Source, Paul-Scherrer Institute Villigen, Switzerland. Phasing was achieved by SHELX and initial automatic model building by ArpWarp ^48, 49^. The orthorhombic crystal form of PcJOS was solved by molecular replacement using PHASER and the model from the tetragonal crystal form with data from beamline ID23_2 at ESRF, Grenoble, France. For PaJOS, data were also collected at beamline X06SA at the Swiss Light Source. The structure was solved by molecular replacement employing a search model predicted by AlphaFold 2 ^35^ and ubiquitin (entry 1UBQ) ^50^. The structure of BpJOS was likewise determined using AlphaFold and PHASER with data from beamline P13 at PETRAIII, EMBL outstation, in Hamburg, Germany. Data collection and refinement statistics is given in Supplementary Tables 2-3.

### Activity-based probe assays

DUBs or UCCs were prediluted to 2× concentration (10⍰µM) in reaction buffer (20⍰mM TRIS pH 7.5, 150⍰mM NaCl and 10⍰mM DTT) and combined 1:1 with 100⍰µM activity-based probes for 18 hours at 20°C. Deviating time points are indicated in the respective legend The reaction was stopped by the addition of 2x Laemmli buffer, and analyzed by SDS-PAGE using Coomassie staining.

### Ubiquitin chain cleavage

DUBs were prediluted in 150⍰mM NaCl, 20⍰mM TRIS pH 7.5 and 10⍰mM DTT. The cleavage was performed at 20°C for the indicated time points with different DUB concentrations (PcJOS/BpJOS: 50 nM, PaJOS: 0.5 µM or as indicated in the respective figure legends) and 25⍰µM diubiquitin (M1, K11, K48, K63 synthesized as described above, K6, K29, K33 purchased from Biomol, K27 from UbiQ) or 20 µM Ub6+ chains (K63; synthesized as described above). The reactions were stopped with 2x Laemmli buffer, resolved by SDS-PAGE, and either Coomassie stained or transferred to PVDF-membranes by Western Blotting.

### Cell culture and Western Blotting

HEK293T cells obtained from ATCC were cultured at 37°C and 5 % CO2 in Dulbecco’s modified eagle medium (DMEM; Gibco) supplemented with 10 % fetal calf serum (FCS) and 1 % Penicillin-Streptomycin. Cells were grown to 90 % confluency before harvest. Collected cell pellets were resuspended in lysis buffer (20 mM Tris pH 7.5, 150 mM NaCl, 0.5 % NP-40, 2 mM EDTA, 5 mM NEM) and sonicated with 3x 10s pulses. Cell debris was cleared by centrifugation at 16,000 x g for 15 min at 4°C. Protein concentration was quantified using a Bradford assay (Roti Quant; Roth) and adjusted to 5 µg/µl total protein content. Unreacted NEM was quenched with 10 mM DTT. Cell lysates were incubated with 10 µM of the corresponding enzyme (BpJOS, BcJOS, PcJOS^70-324^, HeJOS, PaJOS^1880-end^, USP21^196-565^, ATXN3L, JOSD2) for the indicated time. The reaction was quenched by addition of 2x or 5x Laemmli buffer and boiling of the samples at 95°C for 5 min. Samples were resolved on a 12% Tris-Glycine gel and transferred onto PVDF membranes by semi-dry Western Blotting. Membranes were decorated with primary antibodies overnight at 4°C: anti-Ubiquitin (Millipore, 05-944, 1:3000), anti-Diglycyl-Lysine (Lucerna, GX41, 1:500), anti-β-Actin (Santa Cruz, sc-81178, 1:500) or for 1h at RT (anti-DYKDDDDK HRP, Miltenyi Biotech, 1:10000). Secondary HRP-linked antibody was incubated (anti-mouse, Cell Signaling Technology, 1:3000 in 5% milk in PBS-T) for 1h at RT. HRP secondaries were developed using WesternBright chemiluminescent reagent (Advansta, K-12045). Gel images were acquired using ImageLab software 5.2.1.

### Live cell imaging

Time-dependent imaging and analysis of cell death was performed in a IncuCyte® S3 live-cell imager using DRAQ7 as marker. The day prior to transfection 2x10^4^ cells/ well were seeded in 48-well plates. Expression levels of used constructs were checked by preparation of a 6-well plate in parallel for subsequent Western Blotting. The next day, cells were transiently transfected with the following pcDNA5-based plasmids: FLAG-GST^59-712^, FLAG-BpJOS, FLAG-PcJOS^70-324^, FLAG-USP2^258-605^, and MLKL^1-180^-FLAG. If applicable, cells prepared for live cell imaging were treated with 20 µM zVAD-FMK 2h before the measurement. The cell medium was supplemented with 0.1 µM DRAQ7 30 minutes prior to imaging. Cells were imaged at 10x magnification and brightfield and the red channel were set. Cell death scans were performed every 60 minutes for 48 hours. The resulting raw data were processed in Excel by calculating the mean and the standard deviation of the respective replicates.

## Data Availability

All X-ray structures have been deposited at the PDB database under the accession numbers 9F5T (BpJOS), 9FN4 (PcJOS), 9FPA (PcJOS, orthorhombic) and 9G7G (PaJOS). Source data underlying the findings of this study are provided with this article.

## Acknowledgments

We thank Christiane Horst for expert technical assistance. This work was supported by the DFG (Grant HO 3783/3-2 to K.H.) and NWO (VIDI Grant VI. 213.110 to M.P.C.M.). Mass spectrometry was performed at the Proteomics Facility of the CECAD, University of Cologne. Crystals were grown using equipment of the Cologne Crystallization facility (C2f), which is supported by DFG grant INST 216/949-1 FUGG. The synchrotron data were collected at the Swiss Light Source, Paul-Scherrer-Institute, Switzerland, at beamline X06SA, at the European Synchrotron Radiation Facility (ESRF), France at beamline ID23_2 and at PETRAIII, EMBL outstation, in Hamburg, Germany at beamline P13. We thank the staff for their support.

## Author contributions

T.H. and S.K. performed all biochemical and crystallization experiments. M.U. collected the X-ray data and solved the structure, U.B. supervised the crystallography. R.A.d.H. and M.P.C.M designed and synthesized activity-based probes. KH conceived and supervised the project and contributed bioinformatical analyses. All authors contributed to the manuscript.

---

**Supplementary Figure 1. Different cleavage positions within the bacterial Josephin-family. a)** Activity of SnJOS1 and SnJOS2 against K63-linked diubiquitin-VME activity-based probes. **b)** Intact mass spectrometry of mono-ubiquitin cleaved by SnJOS1 (left panel), SnJOS2 (middle panel) and USP21 (right panel). The m/z ratio is depicted on the x-axis, while the deconvoluted masses are shown next to the respective peaks. The input sample shows a single peak, which gets shifted by 114 Da after treatment with SnJOS1 and SnJOS2, but not by USP21 treatment, indicating a removal of the terminal GlyGly motif. **c,d)** Intact mass spectrometry of 6xHis-tagged mono-ubiquitin cleaved by BpJOS (c) or BcJOS (d). The m/z ratio is depicted on the x-axis, while the deconvoluted masses are shown next to the respective peaks. **e**) Intact mass spectrometry of mono-ubiquitin cleaved by PaJOS. The m/z ratio is depicted on the x-axis, while the deconvoluted masses are shown next to the respective peaks. **f)** Intact mass spectrometry of K63-linked diubiquitin cleaved by HeJOS. The m/z ratio is depicted on the x-axis, while the deconvoluted masses are shown next to the respective peaks. The input sample shows a single 17100 Da peak (light blue) corresponding to the monoisotopic mass of diubiquitin, which gets cleaved by HeJOS to a single product (orange) corresponding to the mono-isotopic mass of ubiquitin (8558.6), indicating a conventional cleavage.

**Supplementary Figure 2. Eukaryotic Josephins have no clippase activity. a)** Determination of the cleavage position by activity-based probes. Human ATXN3L and JOSD2 were incubated with Ub^1-75^ or Ub^1-73^-PA probes for 18h. Asterisks (*) mark the shifted bands after the reaction. **b)** Analysis of eucaryotic Josephins cleavage products by Western Blotting. HEK293T cell lysates were incubated with the BpJOS, ATX3NL, JOSD2 or USP21 for 2 h. Substrate deubiquitination and accumulation of mono-ubiquitin is visualized by α-ubiquitin detection (left panel) and accumulation of diGly-remnants is visualized by α-K-ε-GlyGly detection (right panel). α-actin staining serves as a loading control.

**Supplementary Figure 3. Multiple sequence alignment of bacterial Josephin-family members.** Multiple-sequence alignment of all bacterial Josephins covered in this study, in comparison to two human Josephins (bottom lines). Sequences with bold blue names are UCCs, the other ones conventional DUBs. Residues printed on black or grey background are invariant or conservatively replaced in at least 50% of the shown sequences. The active site residues are highlighted in red and the conserved aromatic motif is highlighted in blue.

**Supplementary Figure 4. Comparison of BpJOS and Lb** pro**\*. a)** Activity-based probe reaction of BpJOS and Lb^pro^ against Ub^1-73^-, Nedd8^1-73^ and ISG15^79-154^-PA. **b)** Intact mass spectrometry of ISG15^79-165^ incubated with BpJOS or Lb^pro^*. The m/z ratio is depicted on the x-axis, while the deconvoluted masses are shown next to the respective peaks. The input sample shows a single 9790.1 Da peak (blue) corresponding to the monoisotopic mass of ISG15^79-165^. The mass remains unchanged after incubation with BpJOS(green), while Lb^pro^*(orange) cleaves the terminal GGGTEPGGRS peptide, thereby reducing the mass to 8933.7Da. **c)** Activity of 50 nM BpJOS and 5µM Lb^pro^* against K48 linked diubiquitin.

**Supplementary Figure 5. Crystal structure of PcJOS in complex with linear diubiquitin. a)** Assembly of the PcJOS/diubiquitin complex in the asymmetric unit shown in cartoon representation Linear diubiquitin (lightblue) is bound to the S1 and S1’ sites of one PcJOS molecule (light grey). The proximal ubiquitin is additionally bound to the S1 site of a second PcJOS molecule (dark grey). Active site residues are shown as sticks and coloured orange. **b)** Structural superposition of the two PcJOS molecules shown in a). RMSD: 0.44Å over 1317 atoms. **c)** Activity of wildtype PcJOS and active site mutants (C162A, H284A and D300A) against linear diubiquitin. **d)** Structural superposition of PcJOS (grey) and the human Josephins ATXN3L (green, PDB: 3O65) or JOSD2 (yellow, PDB: 6PGV) shown in cartoon representation. RMSDs: 1.4 Å over 500 atoms (PcJOS/JOSD2) and 3.7 Å over 574 atoms (PcJOS/ATXN3L). **e)** Close-up view on the superposition shown in (d) highlighting active site residues shown as sticks**. f,g)** Activity of PcJOS truncations against a panel of differently linked diubiquitins (f) or activity-based probes (g). **h)** Activity of wildtype PcJOS against linear diubiquitin mutants. E16’A and E18’A refer to the positions within the proximal ubiquitin and correspond to the positions E92A or E94A of the diubiquitin. **i)** Phe-285 stabilizes Leu-73 in the ubiquitin C-terminus. Important residues are shown as sticks and active site residues are coloured orange. Hydrophobic interactions are indicated by yellow, dotted lines. **j)** Activity of wildtype PcJOS or the aromatic motif mutant F285A against linear linked diubiquitin.

**Supplementary Figure 6. Crystal structure of PaJOS. a)** Magnification of the PaJOS active site. Important residues are shown as sticks and coloured green (PaJOS active site) or light blue (ubiquitin C-terminus). **b)** Activity of wildtype PaJOS and active site mutants (C66A, H187A and D203A) against K63-linked diubiquitin. **c)** Structural superposition of PaJOS (green) and BpJOS (orange) shown in cartoon representation. The respective Histidine residues contacting Ile36 of ubiquitin are highlighted as sticks. Hydrophobic interactions are indicated by yellow, dotted lines**d)**. Comparison of Ile-36 patch recognition by PaJOS and human ATXN3L. The superposition is based on the catalytic domains of PaJOS (green) and ATXN3L (grey, PDB: 3O65). The respective ubiquitin molecules are coloured light blue (PaJOS) or lightorange (ATXN3L). **e)** Activity of wildtype PaJOS and aromatic gatekeeper mutant F188A against K63-linked diubiquitin.

**Supplementary Figure 7. Crystal structure of BpJOS. a)** Overview over the asymmetric unit. **b)** Structural superposition of the two BpJOS molecules shown in a). RMSD: 0.2Å over 1308 atoms. **c)** Activity of BpJOS truncations against M1-linked diubiquitin. **d)** Magnification of the BpJOS active site. Important residues are shown as sticks and coloured orange (BpJOS active site) or yellow (ubiquitin C-terminus). The position of the ubiquitin C-terminus in relation to the active site residues supports a cleavage between Arg-74 and Gly-75. **e)** Activity of wildtype BpJOS and active site mutants (C69A, H166A and D182A) against linear diubiquitin. **f)** Comparison of Ile-36 patch recognition by BpJOS and PcJOS. The superposition is based on the catalytic domains of BpJOS (orange) and PcJOS (grey). The respective ubiquitin molecules are coloured yellow (BpJOS) or light blue (PcJOS). **g)** Comparison of RLRGG recognition by PaJOS and BpJOS. The superposition is based on the catalytic domains of PaJOS (green) and BpJOS (orange). The respective ubiquitin molecules are coloured light blue (PaJOS) or yellow (BpJOS). Interactions between the proteases and ubiquitin’s Arg72, Leu73 and Arg74 residues are indicated as yellow dotted lines.

**Supplementary Figure 8. Superposition of bacterial Josephin models generated by AlphaFold.** Structure models of bacterial Josephin candidates (Supplementary Table 1) in complex with ubiquitin were generated using AlphaFold2. In order to predict DUB or UCC-cleavage the catalytic domains were superimposed onto the PaJOS/BpJOS structures. The superimposed structures are shown in cartoon representation and are coloured orange (BpJOS), green (PaJOS) and light purple (candidates). The corresponding ubiquitins are coloured yellow, light blue and pink respectively. RMSDs: 3.7 Å over 136 residues (PtJOS/PaJOS), 3.3 Å over 549 atoms (KrJOS/PaJOS), 1.4 Å over 391 atoms (MtJOS/PaJOS), 3.8 Å over 711 atoms (MxJOS/PaJOS), 1.6 Å over 505 atoms (ScJOS/PaJOS), 1.4 Å over 357 atoms (MlJOS/PaJOS), 2.1 Å over 549 atoms (CsJOS/PaJOS), 2.4 Å over 659 atoms (Pc2JOS/PaJOS).

