## Supplementary figures and images for "A family of bacterial Josephin-like deubiquitinases with an unusual cleavage mode"

### Supplementary Fig 1

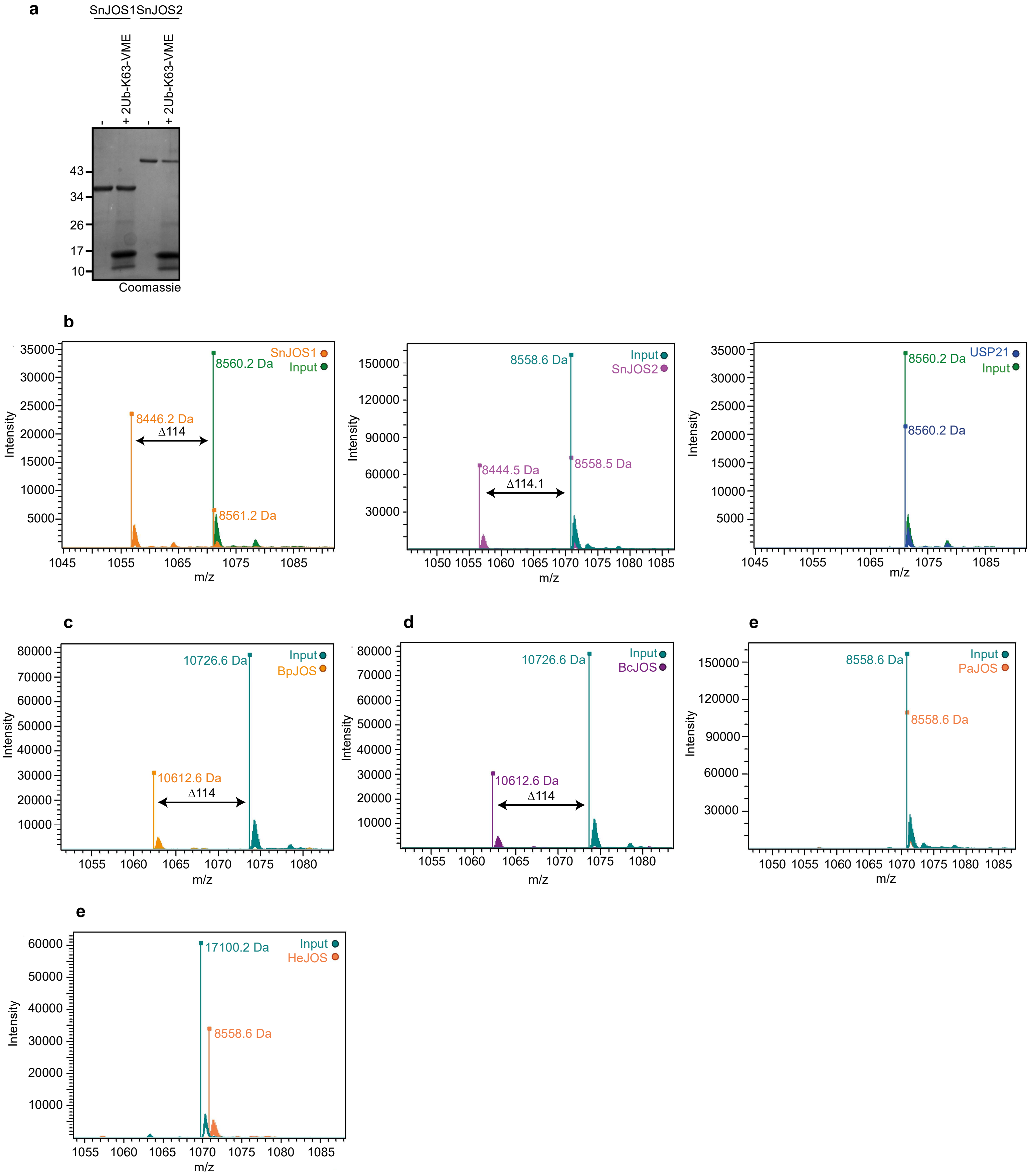


**Supplementary Figure 1**

### Supplementary Fig 2

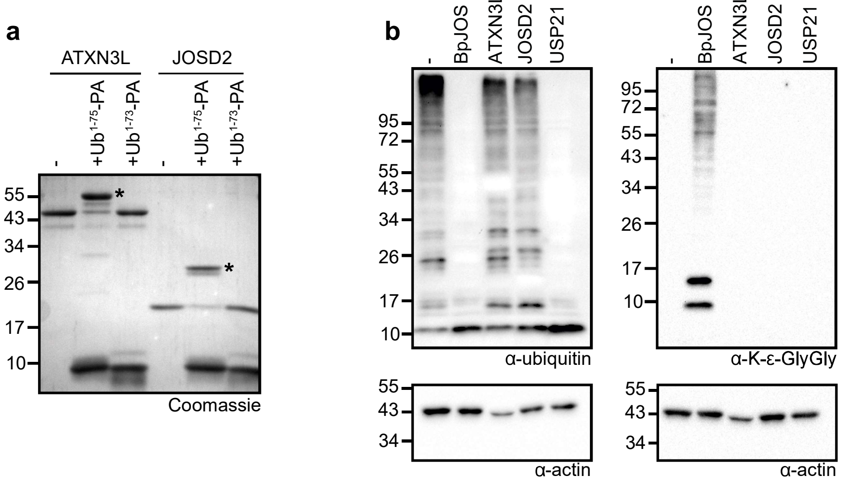


**Supplementary Figure 2**

### Supplementary Fig 4

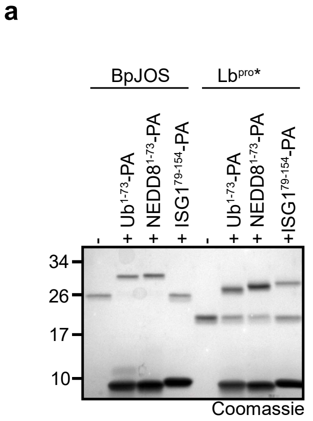


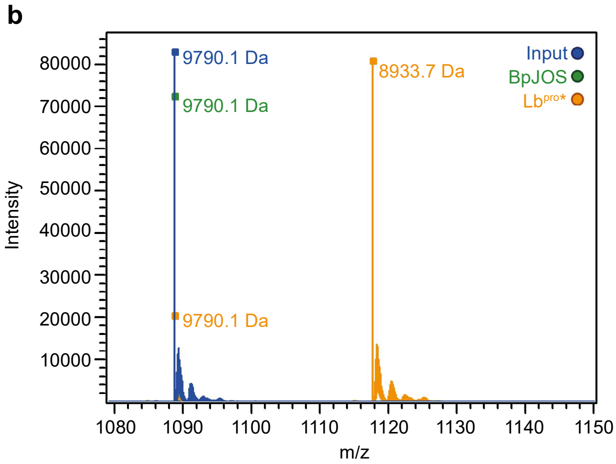


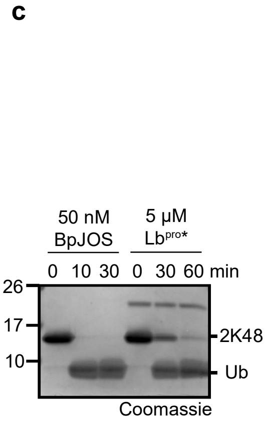


**Supplementary Figure 4**

### Supplementary Fig 5

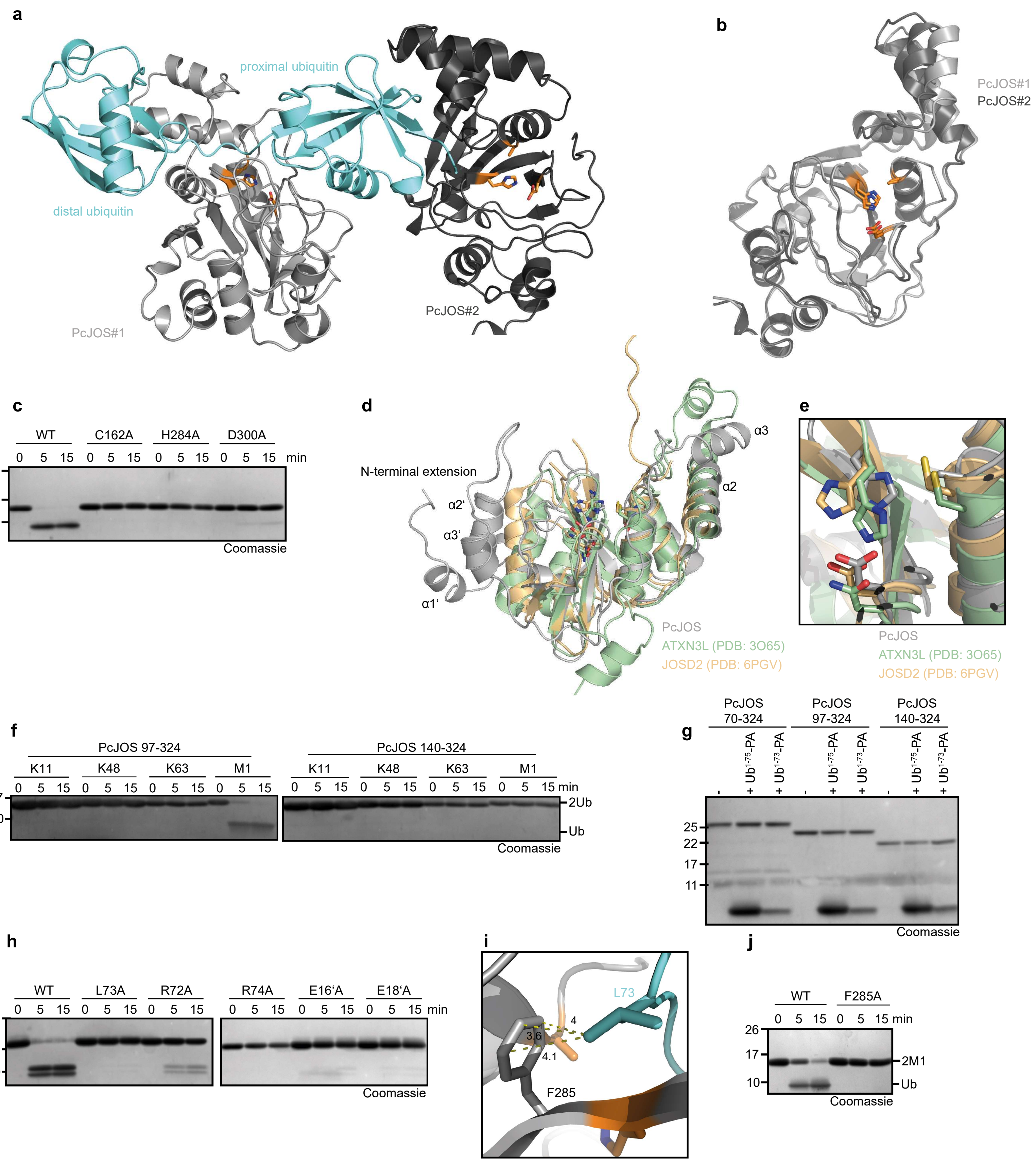


**Supplementary Figure 5**

### Supplementary Fig 6

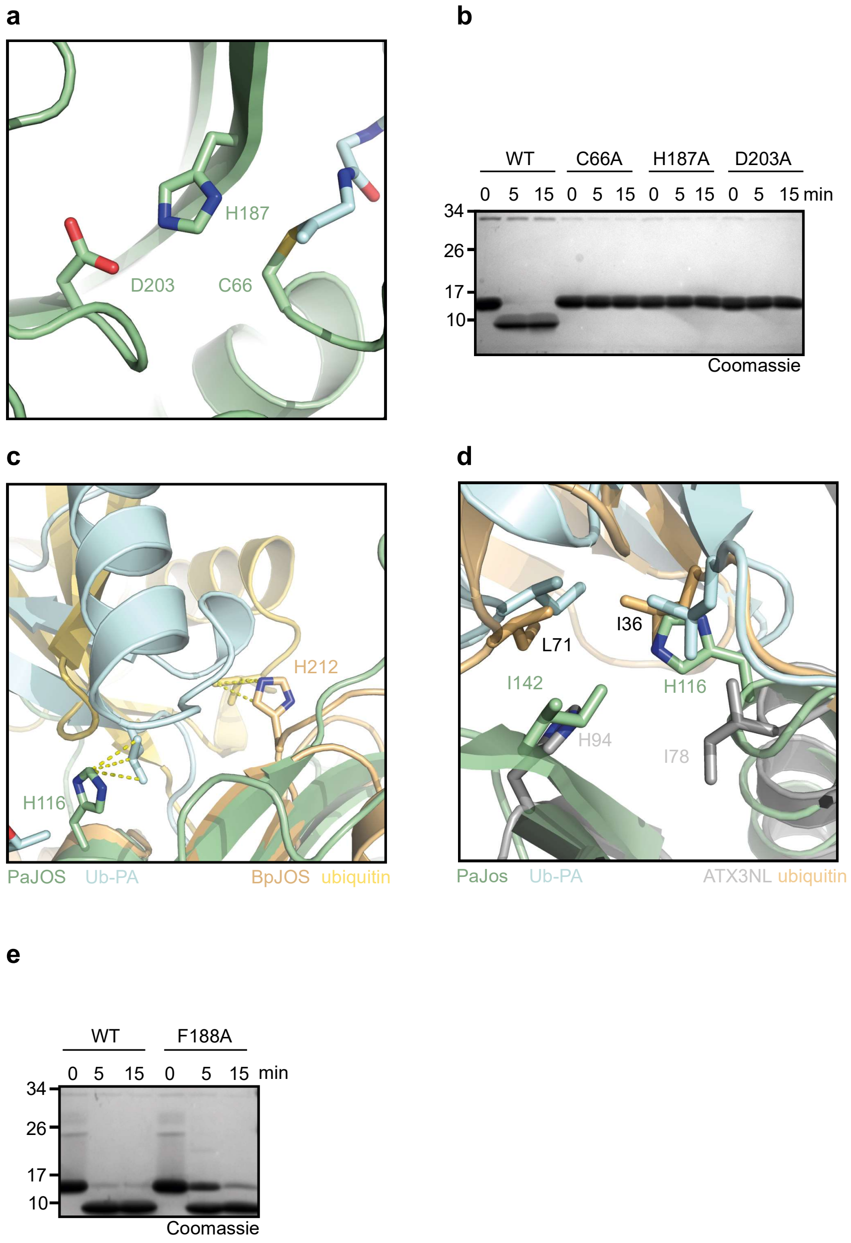


**Supplementary Figure 6**

### Supplementary Fig 7

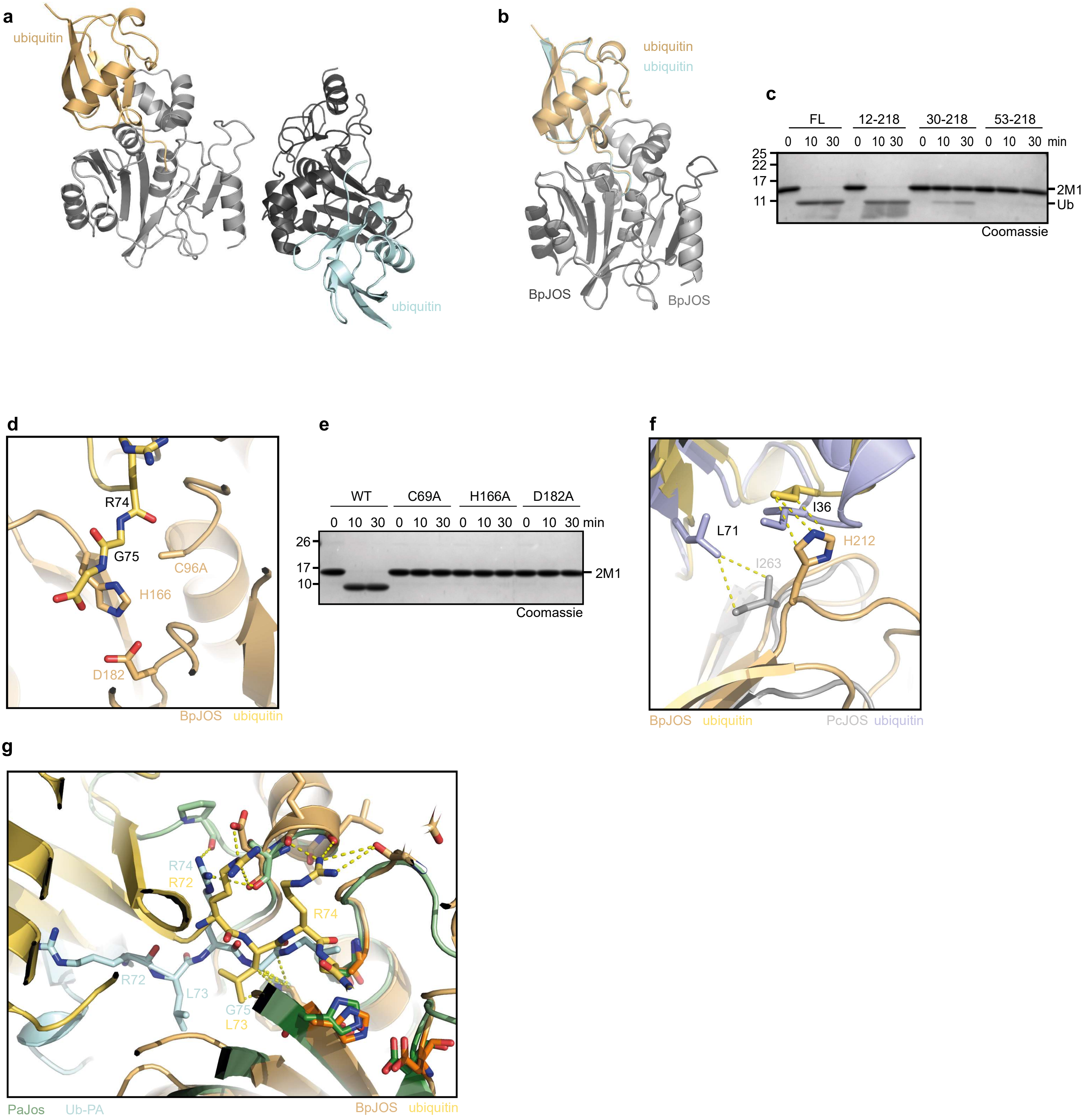


**Supplementary Figure 7**

### Supplementary Fig 8

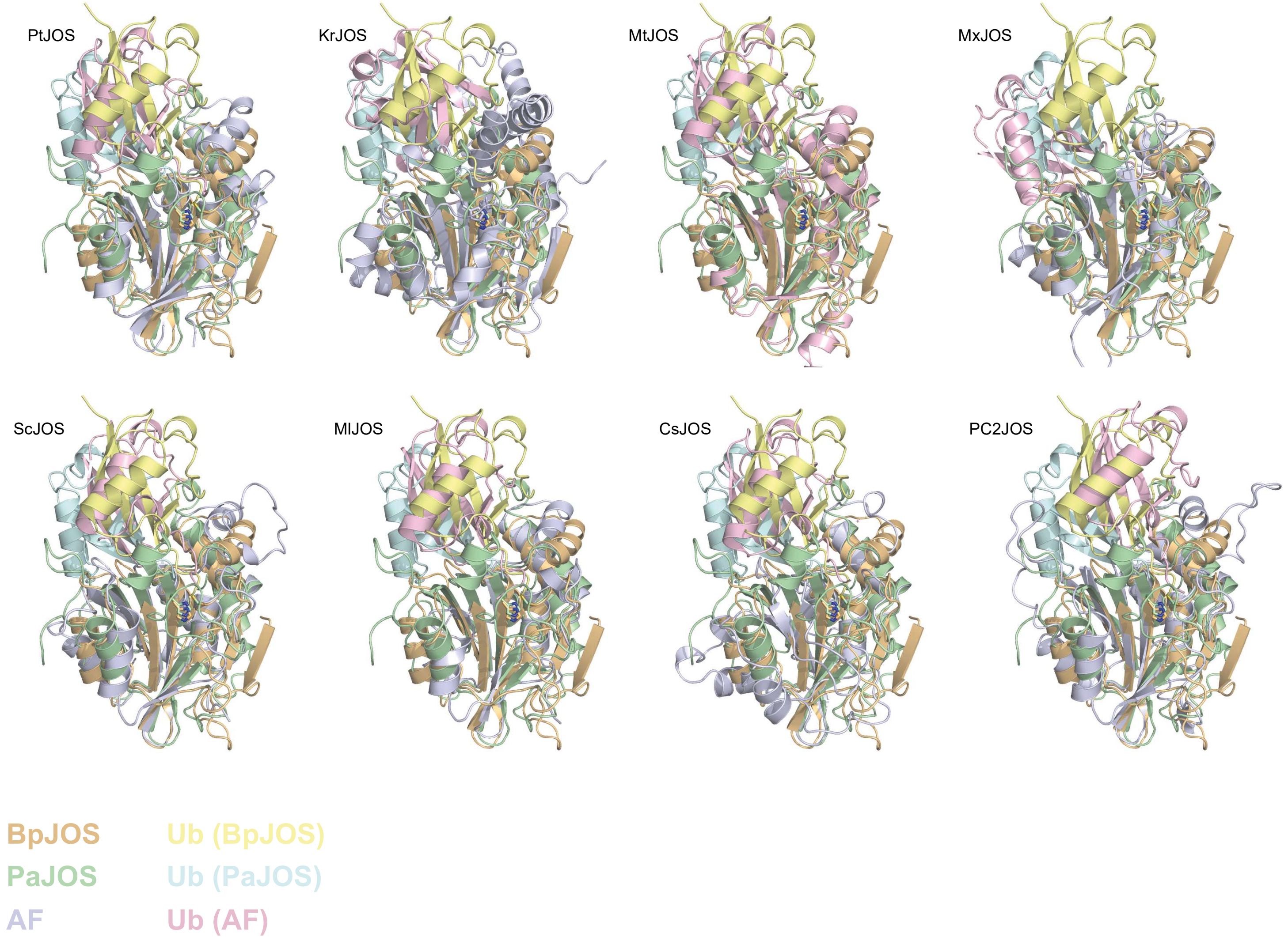


**Supplementary Figure 8**
