## Supplementary Fig 3 for "A family of bacterial Josephin-like deubiquitinases with an unusual cleavage mode"

```
SnJOS1  EKYYCEAQK...........NSFCQMHAANAFLGY.....GAIKPNDLAEYIAKRAA.........KFCHSHLED..................PADGESARASTK.KLLDGVLDIEYGIDLGMVV
SnJOS2  EKYYCEEQK...........NSYCQIHAANAFLGY.....GAIRPSAIAEYVKMRAT.........GFGHDQVEG..................LAMGYGPQDT...KSAEGLLDIENGVDLGMVV
PcJOS   HIAIVNAQE.........TGLGHCGRYAINNALQQ.....EALSQNEFLSLTGQIFS....................................NQLGMSLADV...KNLIQNQD.DFGIDTGVLA
BpJOS   GLFYHEIQR.........VDSGTCAVHAANAYSGS.....SQYNLHHFANAQS........................................NMVGLDYNEA...KGLILQ....DGNDPNFVK
BcJOS   SLFYHEVQH.........RNSGSCGIHAANAYLGE.....PAIDAASVAKECN........................................EIFGADVLSA...DEIAKG.....GADG....
PaJOS   DQFHVEPQV...........ASACAIHSINAFVGG.....PAFDIPTFTTWSTASTA.........AFIGDDAD...ALAPES..........AASGFSPHRV...ERALNLLD...GTPATQGK
HeJOS   PRFYQQFQAAQAALQRGEPSGNMCGALALAHMTGTQPQ.WPQLHRLAMSDLSRHTAP.......................................GRASEAQ...INAARKTV.YDGVYIEHLQ
PtJOS   DFFLKQPHS........GEQKSYCGYYALCHYMGKELD.LNDFRDTVYKQY..........................................VDIGLPEDEI...RQLIAEIG..TGMEGIATG
MtJOS   HKFLPQPNT........QPDCNLCAYYAYYHYTNGGTD.KATFNQRAITHYQ.........................................TRLEMNFADA...QAMYQG.....GNDPSVLE
MxJOS   PPAFLEKQR...........WYQCGLHAVNNLLQR.....PVYAVEDFEAIANEVSAIVP......GIRHK......................ACLGLGCYGA...EVVCAAVA.RQGLCVDTLN
ScJOS   APFHHERQQ........GAASGLCGLHALNAFVGK.....NQFSAENVGERLKGIIAEDQ......GIEAEEALD..QVYPRSVGNLDEHDTTNLVGINPPDM...KTLLTQLG.NEGTIP....
MlJOS   YKFLKQPKT........PPDSKLCGFYALYHFTNGGLT.RDEFIQKATQHYE.........................................TALGLPNKEA...QEMVMD.....GNDPSVLG
KrJOS   SGLYHEMQY.........EASGTCTVHANNHYLASWCE................................REKRPFLPLTPRRLEL.......LFCGIKYRLETEIKEMSKMSA.QCGGQPGSVK
CsJOS   DGWLHEKQE...........PGLCGMHAVNNAIGKIIETKAAFRQKVLTLL..........................................NEIGFSVDAM...PD.....G.EFGVDGEQLR
Pc2JOS  APFHHEAQP........EASSGLCGLHALNAFIGC.....RQFSPRTLGNLVTKAILEGN......QFSQREAEE..ALFPRN.EQTQELNFENFIGLEVPLL...KDVIDEMA.KTHVHP....

ATXN3L  DFIFHEKQE...........GFLCAQHCLNNLLQG.....EYFSPVELASIAHQLDEEERMRMAEGGVTSEEYL.AFLQQPSE..........NMDDTGFFSI...QVISNALK.FWGLEIIHFN
JOSD2   PTVYHERQR...........LELCAVHALNNVLQQ.....QLFSQEAADEICKRLAPDSR..................LNPHR..........SLLGTGNYDV...NVIMAALQ.GLGLAAVWWD

SnJOS1  ......DYLRELESEGVLR..ISVAAIQVGDLS.YSEERGLEFVNHVAGTRHILD...DEFLATK...........SR..AMLGTYQPIHASALRK.....NSDGSWTRIDSSEPSQKVFVD
SnJOS2  ......DFMRHLESEDELKKGVSVTNIEVGELA.YTEENGLEFVNHETQTRHQVD...EEFLAAR...........SR..SILGTFIPVHARALRK.....NTDNSWTEVDSFSPGQAVSKD
PcJOS   ..........QILKQKFNQPV...KESKICDLPDCALSKQQAVENYIGKA........KWVIVAN.IGTEVFDMPSST..HAVYPLTRGHFVALRR.....DADNRWWYLDSRGKNPVNIAL
BpJOS   ..........AVLNESKGAANTAHIAKSKTELADILDHVDRDI...............DRVMVGL.................AGPGESGHWVAFRK.....DGDKKWHKIDSYPRGIRASDP
BcJOS   ..........LPLANAMGAAH.AHKTGDANELKEILGKHQSAI...............DRVMIGI.................SGNDEDAHWVAFRK.....DAGGAWHFINSYPGQNKGSWN
PaJOS   ....DWNIGVSILSPRSGAAM.................ITQVTLPALGDT........DRLIFDVKVGSD......AR..TAAGADDIDHFVAFRK.....DDQGAWWLLDSRSSEVHAPPG
HeJOS   ..........ALLKKNQVAFD.............TLSAIDPADIDRVFTSGRC.....KGVTYGY.................VRPGGTAHFAALRK.....DASGQWWDMDSYHEQPRKIAS
PtJOS   ..................................KYNFIESKDPTPGIDS........GKFMAAT...................NRYSGHWLTYMR.....DKLGKWWEYDSWKNRPTIIGN
MtJOS   ..................................SFQLNLHTTIPTRAVR........KFIVAGT...................MNGGGHFWTIKRI......GWHWWSFDSYHQQAPSRIM
MxJOS   .....CKNTDVALRRLSGPNV.....................................VGLLLNE.VESD......RR..WWQLWRSQRHWLAAVRRATETNSSAYWWIIDSQNCNGRPQKP
ScJOS   ..........ANLANIEVTEV.REVYKDNADLGPVTDQIGDHV...............DRFMVGY...................SGLSKHYVTFRK.....DDDGKWWIIDPQRSRQDSFPD
MlJOS   ..................................LYGLAQSDADTLKKRG........VGVIADT.....................TRGHFFTVRQ.....E.NGIWSSYDSYNHDAAKAYP
KrJOS   ..[32].DKSSALSSNFLTLGVIDIVNQLYSLPKRISSEDKSFKKWETDSKQIREIEQQDSLICTF.............DASEASQQSGHAVCFTKK......NNIWYLQDSNFPEPIQCSP
CsJOS   ..........RILEEGELGIQ..TKTSKISDLPGFKKSKANALAKHVGNS........PWFILYH.NNPEPVPMPRSRKGNATYPIATGHLIAARK.....DNNNQWWMIDSRNPGMVDIPL
Pc2JOS  ..........GIVCETKIINCRGRNGPSNSSLGEPINVVEPSITTSINNV........DRFILGL...................AGIEAHFLTFRK.....DDEGKWWKVDSKSKEQKSFEN
ATXN3L  ......................................NPEYQKLGIDPIN......ERSFICNY......................KQHWFTIRK.....F.GKHWFNLNSLLAGPELISD
JOSD2   ........RRRPLSQLALPQV.....................................LGLILNLPSPVSL........GLLSLPLRRRHWVALRQ.....V.DGVYYNLDSKLRAPEALGD
```

**Supplementary Figure 3**
