## Supplementary Table 1 for "A family of bacterial Josephin-like deubiquitinases with an unusual cleavage mode"

| **Enzyme** | **AccNo (Uniprot)** | **Species** | **Phylum** | **Activity** |
| --- | --- | --- | --- | --- |
| SnJOS1 | F8L436 | *Simkania negevensis* | Chlamydiota | UCC |
| SnJOS2 | F8L650 | *Simkania negevensis* | Chlamydiota | UCC |
| BpJOS | A0A118Q714 | *Burkholderia pyrrocinia* | Pseudomonadota | UCC |
| BcJOS | A0A1J9PQV4 | *Burkholderia catarinensis* | Pseudomonadota | UCC |
| PaJOS | A0A5C0B1L0 | *Pigmenthiphaga aceris* | Pseudomonadota | DUB |
| HeJOS | J3HRE0 | *Herbaspirillum sp.* | Pseudomonadota | DUB |
| PcJOS | A0A2H9SU66 | *Parachlamydia sp.* | Chlamydiota | UCC (M1) |
| PtJOS | A0A0F7LTE5 | *Photorhabdus thracensis* | Pseudomonadota | UCC |
| MtJOS | A0A0V7ZVW0 | *Mastigocoleus testarum* | Cyanobacteriota | UCC |
| MxJOS | A0A7V5I3J3 | hot springs metagenome | n.d. | DUB |
| ScJOS | A0A8A4TFZ2 | *Sulfidibacter corralicola* | Acidobacteriota | UCC |
| MlJOS | A0A540WM54 | *Myxococcus llanfairpwllgwyngyllgogerych-wyrndrobwllllantysiliogogogochensis* | Myxococcota | UCC |
| KrJOS | A0A0L0ANS7 | *Klebsiella sp RIT-PI-d* | Pseudomonadota | - |
| CsJOS | KAF3361746^*^ | *Chlamydiales bacterium STE3* | Chlamydiota | UCC (M1) |
| Pc2JOS | MCE5317260^*^ | *Parachlamydia sp* | Chlamydiota | UCC |

*RefSeq database accession

**Supplementary Table 1. Enzymes characterized in this study.**
