## Supplementary Table 2 for "A family of bacterial Josephin-like deubiquitinases with an unusual cleavage mode"

**Supplementary Table 2. Data collection and refinement statistics.**

|  | PcJOS - orthorhombic | PcJOS - tetragonal |
| --- | --- | --- |
| Wavelength | 0.8731 | 0.978 |
| Resolution range | 43.9 - 2.18 (2.23 - 2.18) | 57.76 - 2.151 (2.18 - 2.15) |
| Space group | P 21 21 21 | P 41 21 2 |
| Unit cell | 54.22 108.18 149.61 90 90 90 | 88.765 88.765 147.545 90 90 90 |
| Total reflections | 360659 (17316) | 435289 (19405) |
| Unique reflections | 88394 (5383) | 61147 (2762) |
| Multiplicity | 4.1 (3.2) | 7.1 (7.0) |
| Completeness (%) | 99.88 (99.23) | 99.88 (98.42) |
| Mean I/sigma(I) | 6.44 (0.88) | 10.13 (0.75) |
| Wilson B-factor | 39.00 | 45.02 |
| R-merge | 0.1295 (1.3) | 0.1274 (2.393) |
| R-meas | 0.1493 (1.538) | 0.1375 (2.583) |
| R-pim | 0.07319 (0.8028) | 0.05119 (0.9636) |
| CC1/2 | 0.995 (0.369) | 0.998 (0.332) |
| CC* | 0.999 (0.734) | 1 (0.706) |
| Reflections used in refinement | 46730 (2833) | 32753 (1445) |
| Reflections used for R-free | 2263 (136) | 1635 (72) |
| R-work | 0.1850 (0.2810) | 0.1867 (0.3628) |
| R-free | 0.2128 (0.3200) | 0.2045 (0.3661) |
| Number of non-hydrogen atoms | 4940 | 2522 |
| macromolecules | 4684 | 2389 |
| ligands | 26 | 10 |
| solvent | 230 | 123 |
| Protein residues | 599 | 306 |
| RMS(bonds) | 0.006 | 0.002 |
| RMS(angles) | 0.81 | 0.58 |
| Ramachandran favored (%) | 97.29 | 98.34 |
| Ramachandran allowed (%) | 2.54 | 1.66 |
| Ramachandran outliers (%) | 0.17 | 0.00 |
| Rotamer outliers (%) | 0.00 | 0.38 |
| Clashscore | 1.27 | 0.83 |
| Average B-factor | 53.14 | 49.08 |
| macromolecules | 53.41 | 49.08 |
| ligands | 76.14 | 67.00 |
| solvent | 45.03 | 47.64 |

Statistics for the highest-resolution shell are shown in parentheses
