## Supplementary Table 3 for "A family of bacterial Josephin-like deubiquitinases with an unusual cleavage mode"

**Supplementary Table 3. Data collection and refinement statistics.**

|  | PaJOS/Ub-PA complex | BpJOS/2Ub complex |
| --- | --- | --- |
| Wavelength | 1.000 | 0.9793 |
| Resolution range | 49.03 - 1.89 (1.93 - 1.89) | 53.53 - 2.561 (2.652 - 2.561) |
| Space group | P 1 21 1 | P 1 21 1 |
| Unit cell | 131.946 81.316 167.392 90 90.23 90 | 89.61 35.51 117.64 90 102.205 90 |
| Total reflections | 1792663 (61130) | 63680 (2833) |
| Unique reflections | 271313 (14118) | 22646 (1442) |
| Multiplicity | 6.6 (4.3) | 2.8 (2.0) |
| Completeness (%) | 95.61 (73.50) | 94.22 (61.97) |
| Mean I/sigma(I) | 10.51 (0.62) | 10.43 (1.05) |
| Wilson B-factor | 39.34 | 56.97 |
| R-merge | 0.072 (1.77) | 0.07414 (0.7163) |
| R-meas | 0.078 (2.01) | 0.09137 (0.9389) |
| R-pim | 0.030 (0.944) | 0.05262 (0.5996) |
| CC1/2 | 0.999 (0.369) | 0.997 (0.542) |
| CC* | 1.00 (0.734) | 0.999 (0.838) |
| Reflections used in refinement | 270643 (13788) | 22638 (1442) |
| Reflections used for R-free | 2116 (112) | 1998 (127) |
| R-work | 0.206 (0.4150) | 0.2097 (0.3573) |
| R-free | 0.2303(0.4439) | 0.2527 (0.4402) |
| Number of non-hydrogen atoms | 29032 | 0.950 (0.671) |
| macromolecules | 28227 | 0.927 (0.517) |
| ligands | 48 | 4456 |
| solvent | 757 | 4442 |
| Protein residues | 3705 | 0 |
| RMS(bonds) | 0.005 | 14 |
| RMS(angles) | 0.80 | 561 |
| Ramachandran favored (%) | 98.38 | 0.003 |
| Ramachandran allowed (%) | 1.59 | 0.56 |
| Ramachandran outliers (%) | 0.03 | 96.56 |
| Rotamer outliers (%) | 0.43 | 3.44 |
| Clashscore | 2.49 | 0.00 |
| Average B-factor | 51.30 | 0.63 |
| macromolecules | 51.51 | 2.27 |
| ligands | 39.29 | 64.58 |
| solvent | 44.12 | 64.64 |

Statistics for the highest-resolution shell are shown in parentheses
